# An exploration of metabolite and gene responses in mouse skeletal muscles responding to acute sedentary hypoxia

**DOI:** 10.1101/2021.06.03.446848

**Authors:** Feng Xue, Gang Huang, Xi Wang, Jie Deng, Lingxia Pang, Zhuohui Gan

**Author notes:** **Correspondence**, Zhuohui Gan, The School of Basic Medical Sciences, Wenzhou Medical University, Ouhai District, Wenzhou, Zhejiang, 325035, China.

## Abstract

Skeletal muscles are involved in responses to acute hypoxia as the largest organ in the body. However, as a hypoxic-tolerant tissue, responses in skeletal muscles caused by acute sedentary hypoxia are much less studied. We measured metabolites in skeletal muscles from mice exposed to 8% O_2_ for 0 minute, 15 minutes and 2 hours and studied the potential relationship between metabolite levels and mRNA levels by reconstructing genome-based metabolic networks and meta-analyzing differentially expressed genes acquired in skeletal muscles after 2 hours of 8% O_2_ exposure. The metabolite measurement indicated a significant increase in glutamine metabolism but not lactate metabolism in mouse skeletal muscles after 2 hours of hypoxia, where the metabolic responses as a whole were moderate. The central-dogma based metabolic flux analysis suggested an involvement of glutamine metabolism, though, as a whole, metabolite changes and gene changes didn’t show a high correlation. Among metaoblites, glutamine metabolism indicated a significant response and a consistent change which could be interpreted by genome-based network analysis. In summary, though this study suggested a moderate metabolic response which has a weak correlation with gene expression change as a whole, glutamine metabolism indicated rapid responses in skeletal muscles responding to acute sedentary hypoxia.

## Background

Hypoxia is widely studied as a common clinical symptom related to many diseases such as pulmonary obstruction which causes systemic acute hypoxia [1–4]. Skeletal muscles are involved in hypoxic responses as the largest organ in the body which make up to ~45% to 55% of body mass [2].

During hypoxia induced by exercise, skeletal muscles as the major responsive tissue are studied widely. A number of changes such as increases in lactate, acetylcarnitine, TCA cycle and decreased glycogenolysis are revealed [5–8]. However, as to acute sedentary hypoxia, hypoxia-sensitive tissues such as brain and heart are studied more frequently, while studies in skeletal muscles, a hypoxia-tolerant tissue, are lagged [5, 9–14].

By a transcriptome-wide investigation, we revealed a number of gene expression changes in skeletal muscles from mice exposed to 8% O_2_ for 2 hours [11]. To further our understanding of changes in skeletal muscles during an acute sedentary hypoxia, in this study, we investigated metabolites in skeletal muscles from mice exposed to 8% O_2_ for 0 minute, 15 minutes and 2 hours. Furthermore, we evaluated the potential relationship between changes on gene expression and that of metabolites by predicting metabolic changes using a genome-based metabolic flux network analysis. The results suggest a moderate metabolic change generally, while glutamic acid indicated a significant increase after 2-hour of 8% O_2_ exposure in mouse skeletal muscles. The prediction of metabolic responses using genome-based metabolic network analysis which was reconstructed based on central dogma weren’t fully reflected by measured metabolites, suggesting central dogma might not fully cover the relationship among RNA, protein and metabolite for situations such as acute hypoxia. Among measured metabolites, glutamine metabolism indicated a significant increase which was predicted by the genome-based metabolic network analysis, suggesting a role of glutamine metabolism of skeletal muscles responding to acute sedentary hypoxia.

## Methods

### Animals

Male 3-month-old wild-type C57BL/6 mice weighing 25-30 g were housed with a standard diet and adapted to the environment for at least 24 hours before experiments.

### Hypoxic exposure and sample collection

The exposure protocol was the same as described in Gan et al. [11]. Briefly, at the experimental day, mice started to fast at 9:00 a.m.. All mice were fasted for at least 4 hours before sample collection. Mice were exposed to room air (control) or normobaric 8% O_2_ (8% O_2_, 92% N_2_) for 15 minutes (15mins) or 2 hours (2hrs) in individual tubes ventilated at 200 mL/min. At the end of exposure, mice were immediately euthanized by cervical dislocation to minimize metabolic disturbances in skeletal muscles. Hind limbs were cut and flash frozen in liquid nitrogen to minimize potential effects on metabolic state. Plantaris muscles, which are relatively balanced in fast glycolytic fibers and slow oxidative fibers [15], were then dissected from thawed frozen hind limbs on ice in 3-5 minutes [11, 16]. The dissected skeletal muscles were stored in liquid nitrogen or −80°C before further processing.

### Metabolite measurement

Plantaris muscles (25~30 mg) were cut into halves for RNA extraction and metabolite measurement respectively. The small half plantaris muscles (5~10 mg) were homogenized in 1 mL ice-cold 80% methanol containing stable-isotope labelled internal standards. The vortex-mixed homogenate was incubated for 30 minutes at −20°C, centrifuged at 17000 ×g for 15 minutes at 4°C. The protein pellet was redissolved in 1 mL of 0.1 M NaOH solution. The protein concentration of protein solution was measured using the Lowry method and used for the normalization of metabolite concentration among samples. The supernatant was moved to a clean tube and dried in a centrifugal evaporator (SPD121P Speed Vac concentrator, Thermo Fisher) at 36°C and reconstituted in 100 μL solution containing 10% methanol and 0.1% formic acid by vortexing, orbital shaking and sonication. The solution was then analyzed by a liquid chromatography-electrospray tandem mass spectrometry (Sciex API4000) [17]. Briefly, authentic standards were infused and monitored by scheduling them at the corresponding retention times. 5μL sample solution was injected in the analytical system (CTC Analytics HTC PAL autosampler) and separated by a reversed-phase HPLC column (Mac-mod analytical) with a 0.3 mL/min flow rate at 25°C, by means of a simple binary acetonitrile (B) partioning in water gradient, both containing 0.1% formic acid. Positive ionization gradient was 5%B for the first 3 minutes, followed by 40%B for 16 minutes, 100%B for 26 minutes and 0%B for 2 minutes. MultiQuant software (AB Sciex) was used for peak integration and quantification. Metabolite concentrations were calculated using the authentic standard in six to eight non-zero levels calibration curves within 85-115% back-calculated accuracies from nominal spanning physiological range concentrations, with 1/x of concentration weight, to compensate for different variance at low concentration, and coefficients of correlation, 0.99, or higher. Since the quantity of plantaris samples for metabolite measurement (5~10 mg) was small, we didn’t match the mass for each sample for further normalization of metabolite concentration, instead, metabolite concentrations were normalized using its protein concentration. The sample size was relatively small (n=3 for each group) for metabolite analysis since we matched samples for both metabolite measurement and RNA measurements. Thus, metabolites whose standard derivation was greater than 50% of the group average in all three groups or whose normalized concentration was less than 0.05 were excluded from further analysis. Compared with the control, metabolites with p <0.05 were considered as significantly changed. Lactate levels in plantaris were measured as previously described [16]. Standards and other chemical reagents were purchased from Sigma.

### RNA sequencing and RT-PCR

RNAs of plantaris from mice exposed to 8% O_2_ for 2 hours were extracted and sequenced as described [11]. Briefly, RNA was extracted using Qiagen RNeasy fibrous tissue mini kit (#74704, Qiagen). Libraries of extracted RNAs whose A260/A280 >1.8 and RIN >8 were built using the Illumina TruSeq^TM^ RNA sample preparation kit (RS-122-2001, Illumina) and sequenced by an illumina HiSeq 2000 system with paired-end 100-bp reads. The RNA-seq raw data were available in NIH National Center for Biotechnology Information GEO database as GSE81286. To check RNA levels using RT-PCR, cDNAs were synthesized using HiScript II Q RT SuperMix (#R223-01, Vazyme). RT-PCR measurements were conducted using a StepOnePlus Real-Time PCR System (Thermo Scientific) with ChamQ™ Universal SYBR qPCR Master Mix (#Q711-02/03, Vazyme) following manufacturer’s guidelines. Primers for mRNAs (Sangon) are listed in Table S1.

### RNA-seq alignment, assembly and differentially expressed gene identification

To acquire a robust and conserved set of differentially expressed genes (DEGs), we aligned and assembled RNA-seq raw data to UCSC.mm10 or mm9 using TopHat2/Bowtie2 or Hisat2/Stringtie respectively. Several tools including AltAnalyze (limma-based), Cuffdiff or DESeq2 which developed different algorithms to identify DEGs were applied to identify DEGs respectively [18–21]. The threshold of differentially expressed genes was set at p<0.01 for AltAnalyze, adj.p<0.01 for DESeq2 and q value<0.01 for Cuffdiff, respectively. The overlapped genes among three DEG sets were defined as conserved differentially expressed genes (cDEGs). The cDEGs were further analyzed using ToppFun for functional annotation clustering or RAPHGA for pathway analysis [11, 22]. The analyzed results were visualized by R package ‘GOplot’ [23]. GO clusters with a Bonferroni q<0.05 and mapped genes>3 were kept for further visualization. The GO clusters or pathways with an overlap rate>0.7 were removed before its visualization.

### Metabolic modeling by genome-based metabolic network reconstruction

To study the potential effect of gene expression changes on metabolism, differentially expressed genes were first mapped to iMM1415 model, a model describing a metabolic flux network which contains 1415 mouse metabolic related genes and 3727 metabolic reactions, to identify metabolic related genes [24, 25]. The distribution of mapped genes in the metabolic network was visualized by Escher [24]. Then, we evaluated the effects of mapped genes’ changes on metabolic network using MetPath, a method we developed for genome-based metabolic analysis [25, 26]. Briefly, based on the global metabolic reconstruction of mouse metabolism [25], a skeletal muscle-specific metabolic network was generated using published gene expression datasets for skeletal muscles and an established model construction algorithm called mCADRE [27, 28]. The differentially expressed genes along with their folds were then mapped to the established skeletal muscle-specific metabolic network. The metabolic reactions involving the mapped genes were analyzed and the potential effects were evaluated quantitatively [26]. This analysis was conducted in MATLAB.

### Statistical analysis

All data presented were means±s.d.. Metabolite data were analyzed by unpaired twotailed Student’s t-test and considered significant if p<0.05. These statistical tests were performed by Excel or R. The sample sizes of each group were described in the Results section.

## Results

### Metabolic responses in skeletal muscles from mice exposed to normoxia or acute hypoxia

Though blood lactate increased quickly in 15 minutes after mice were exposed to 8% O_2_, the change of lactate in skeletal muscles was not significant with lactate levels at 0.93±0.13, 1.15±0.21 and 0.71±0.22 mM for normoxia, 15 minutes and 2 hours of hypoxia respectively (Fig. 1).

**Fig. 1.**
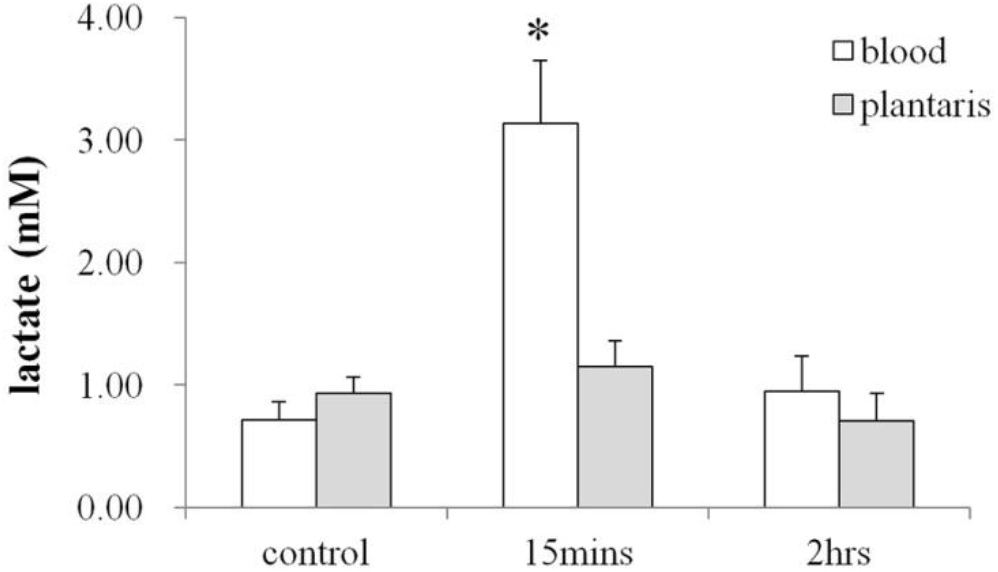
The level of lactate in plantaris or blood. Plantaris muscles were collected from mice exposed to 8% O_2_ for 0 minute (control, n=6), 15 minutes (15mins, n=3) or 2 hours (2hrs, n=6). Blood lactate was previously measured in mice exposed to 8% O_2_ for 0 minute (control, n=5), 15 minutes (15mins, n=5) or 2 hours (2hrs, n=6). All values are mean±s.d.. *: different from that of control samples, p<0.05

Using a LC-MS/MS system, we scanned 132 metabolites totally. Metabolites with big variances (s.d.>50%) or very low levels (normalized concentration<0.05) were excluded for further analysis. The 38 metabolites for further analysis were listed in Table 1. Most of these metabolites in plantaris didn’t change in 15 minutes of hypoxia. In 2 hours of hypoxia, a significant increase in glutamic acid was observed, where glutamic acid was increased by 96%, from 22.05±4.04 to 43.27±6.35 (Fig. 2). Both cytidine and uridine were decreased in skeletal muscles after 2 hours of hypoxic exposure with levels from 0.35±0.06 to 0.18±0.04 and 0.23±0.09 to 0.06±0.05 respectively.

**Fig. 2.**
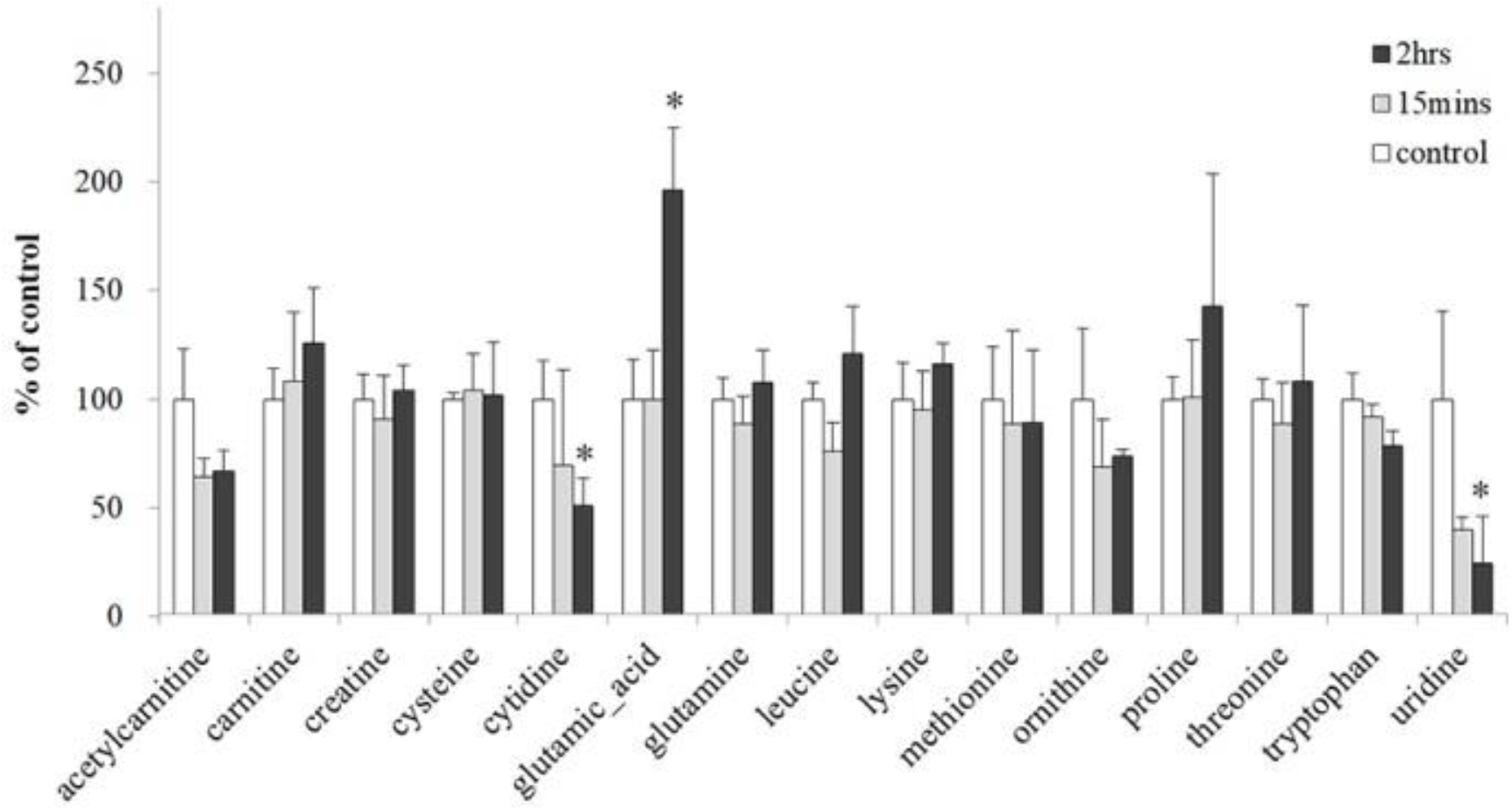
The level of metabolites in plantaris. Plantaris samples were collected from mice exposed to 8% O_2_ for 0 minute (n=3), 15 minutes (n=3) or 2 hours (n=3), respectively. All values are mean±s.d.. *: different from that of control, p<0.05

**Table 1.** Metabolite levels in mouse plantaris. Plantaris samples were collected from mice exposed to normoxia (control, n=3), 8% O_2_ for 15 minutes (15mins, n=3) and 2 hours (2hrs, n=3).

|  | control |  | 15mins |  |  | 2hrs |  |  |
| --- | --- | --- | --- | --- | --- | --- | --- | --- |
|  | mean | SD | mean | s.d. | p | mean | s.d. | p |
| [allo]+isoleucine | 0.25 | 0.05 | 0.23 | 0.08 | 0.78 | 0.33 | 0.09 | 0.21 |
| 1-methylhistidine | 0.75 | 0.02 | 0.74 | 0.10 | 0.80 | 0.86 | 0.10 | 0.16 |
| 5-aminolevulinic | 0.09 | 0.02 | 0.10 | 0.03 | 0.66 | 0.11 | 0.04 | 0.57 |
| 5-methylhistidine | 3.08 | 0.42 | 3.03 | 0.70 | 0.91 | 2.67 | 0.76 | 0.46 |
| acetylcarnitine | 3.47 | 0.80 | 2.23 | 0.30 | 0.06 | 2.32 | 0.33 | 0.08 |
| alanine+sarcosine | 35.13 | 9.79 | 26.68 | 9.87 | 0.35 | 28.39 | 8.24 | 0.41 |
| allo+delta-hydroxylysine | 0.88 | 0.03 | 0.86 | 0.11 | 0.77 | 0.96 | 0.06 | 0.14 |
| alpha-aminoadipic_acid | 0.28 | 0.04 | 0.27 | 0.05 | 0.80 | 0.37 | 0.05 | 0.08 |
| arginine | 3.21 | 1.36 | 1.78 | 0.33 | 0.15 | 2.94 | 1.93 | 0.85 |
| argininosuccinic | 0.10 | 0.01 | 0.10 | 0.01 | 0.83 | 0.11 | 0.01 | 0.14 |
| beta-alanine | 0.75 | 0.24 | 0.73 | 0.21 | 0.91 | 0.68 | 0.32 | 0.76 |
| carnitine | 9.99 | 1.43 | 10.84 | 3.13 | 0.69 | 12.58 | 2.55 | 0.20 |
| carnosine | 8.67 | 0.53 | 8.07 | 0.93 | 0.39 | 6.39 | 1.86 | 0.11 |
| citrulline | 2.28 | 0.62 | 2.50 | 0.47 | 0.65 | 2.95 | 1.05 | 0.40 |
| creatine | 483.38 | 55.58 | 439.96 | 95.50 | 0.53 | 502.51 | 56.43 | 0.70 |
| creatinine | 29.05 | 10.99 | 37.23 | 28.90 | 0.67 | 41.34 | 24.02 | 0.47 |
| cysteine | 207.33 | 6.07 | 215.38 | 35.20 | 0.72 | 211.54 | 50.49 | 0.89 |
| cytidine | 0.35 | 0.06 | 0.24 | 0.15 | 0.33 | 0.18 | 0.04 | 0.02 |
| glutamic_acid | 22.05 | 4.04 | 21.99 | 5.08 | 0.99 | 43.27 | 6.35 | 0.01 |
| glutamine | 21.18 | 2.06 | 18.82 | 2.71 | 0.30 | 22.82 | 3.12 | 0.49 |
| glycine | 58.40 | 10.23 | 59.38 | 28.27 | 0.96 | 60.01 | 29.10 | 0.93 |
| histidine | 3.22 | 0.79 | 3.19 | 0.70 | 0.97 | 3.36 | 0.73 | 0.83 |
| leucine | 1.78 | 0.14 | 1.35 | 0.24 | 0.06 | 2.16 | 0.39 | 0.19 |
| lysine | 12.59 | 2.10 | 11.98 | 2.23 | 0.75 | 14.61 | 1.23 | 0.22 |
| methionine | 0.84 | 0.20 | 0.75 | 0.36 | 0.72 | 0.75 | 0.28 | 0.68 |
| ornithine | 0.67 | 0.22 | 0.46 | 0.14 | 0.25 | 0.50 | 0.02 | 0.24 |
| phenylalanine | 0.30 | 0.05 | 0.34 | 0.07 | 0.48 | 0.42 | 0.15 | 0.29 |
| pipecolic | 0.21 | 0.07 | 0.45 | 0.43 | 0.39 | 0.62 | 0.39 | 0.15 |
| proline | 2.63 | 0.28 | 2.66 | 0.69 | 0.95 | 3.75 | 1.61 | 0.30 |
| serine | 4.41 | 1.35 | 8.66 | 5.52 | 0.27 | 4.67 | 2.18 | 0.87 |
| serotonin | 0.16 | 0.02 | 0.17 | 0.07 | 0.80 | 0.17 | 0.01 | 0.39 |
| taurine | 277.07 | 13.71 | 229.38 | 126.26 | 0.55 | 305.17 | 37.14 | 0.29 |
| threonine | 3.83 | 0.36 | 3.40 | 0.73 | 0.42 | 4.14 | 1.34 | 0.72 |
| trimethylamine-N-oxide | 0.41 | 0.08 | 0.48 | 0.09 | 0.42 | 0.40 | 0.13 | 0.91 |
| tryptophan | 0.63 | 0.07 | 0.58 | 0.04 | 0.36 | 0.50 | 0.04 | 0.06 |
| tyrosine | 0.59 | 0.16 | 0.51 | 0.33 | 0.74 | 0.59 | 0.30 | 0.99 |
| uridine | 0.23 | 0.09 | 0.09 | 0.01 | 0.06 | 0.06 | 0.05 | 0.05 |
| valine | 0.63 | 0.08 | 0.87 | 0.36 | 0.31 | 0.68 | 0.10 | 0.54 |
| protein(mg/mL) | 3.54 | 0.15 | 3.62 | 0.59 | 0.83 | 3.25 | 0.27 | 0.17 |

### Gene expression of skeletal muscles from mice exposed to normoxia or 2 hours of hypoxia

Using DESeq2, AltAnalyze and Cuffdiff, we identified 825, 767 and 1057 DEGs respectively in plantaris from mice exposed to 8% O_2_ for 2 hours. The venn-diagram of DEGs identified different tools was shown in Fig. 3. This result suggests that analysis method could significantly affect the identification of DEGs. About 350 DEGs were conserved among three sets as shown in Table S2. The fold changes calculated by different tools were comparable. Though differences existed in identified DEGs, pathway analysis results indicated a high similarity among three sets as shown in Table S2, suggesting that the key DEGs were conserved among these three DEG sets. Thus, we used the conserved DEGs (cDEGs) for further enrichment analyses.

**Fig. 3.**
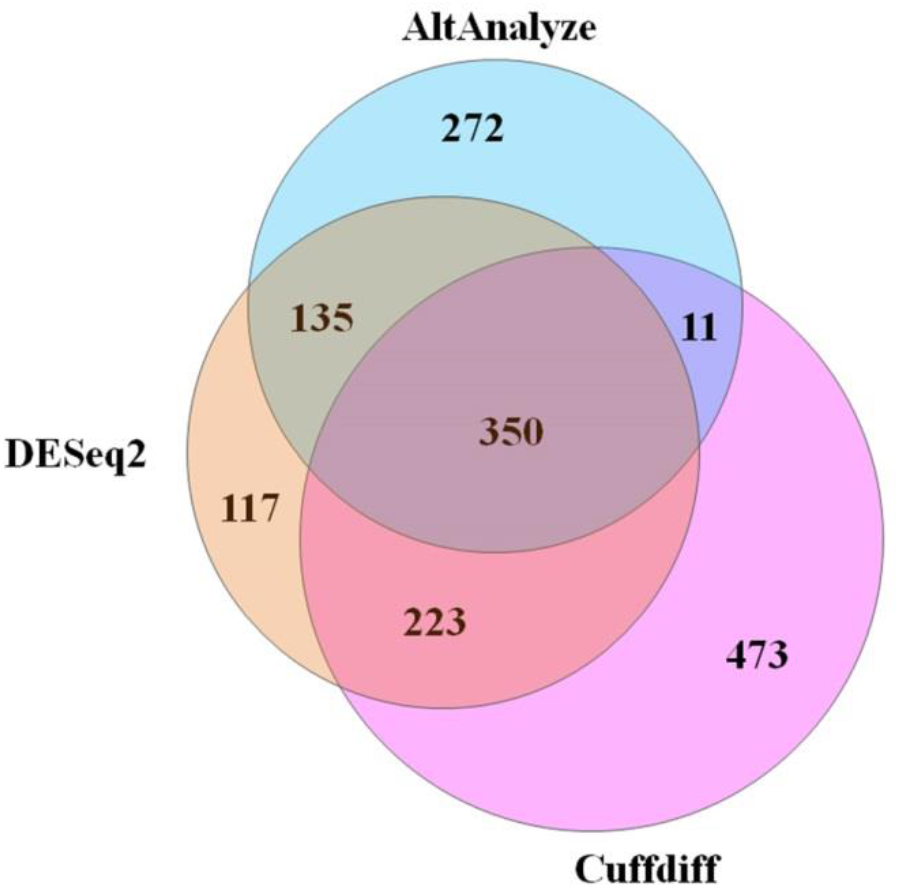
Venn diagram of differentially expressed genes identified by DESeq2, AltAnalyze or Cuffdiff from RNA-seq data of skeletal muscles from mice exposed to normoxia (n=3) or 2 hours of 8% O_2_ (n=3). The radius of each circle is proportional to its total number of DEGs

Pathway analysis identified 47 cDEG-enriched pathways where mmu04512 ‘ECM-receptor interaction’ and mmu04510 ‘focal adhesion’ were the most significant as shown in Table S3. GOplot removed 16 pathways whose overlap rate was greater than 0.7, the remaining pathways were shown in Fig. 4A. Most pathways showed a downregulation at the gene expression level, though it might not mean a down-regulation at the protein or function level. Signaling pathways such as PI3K-Akt signaling pathway, FoxO signaling pathway were highlighted in the pathway analysis results. It is interesting that though down-regulated genes took only two-thirds of mapped genes, they explained most of cDEG-enriched pathways. The only up-regulated cDEG enriched pathway was cytochrome P450 metabolism which contained cDEGs encoding glutathione s-transferase, sulfotransferase and aldehyde dehydrogenase. The functional annotation clustering analysis results as shown in Table S4 also indicated a strong and consistent down-regulation in collagen genes as shown in Fig. 4B, which was confirmed by a RT-PCR measurement (Fig. 5). A number of binding-relevant terms such as enzyme-binding, growth factor-binding were highlighted in the functional clustering results as shown in Fig. 4C.

**Fig. 4.**
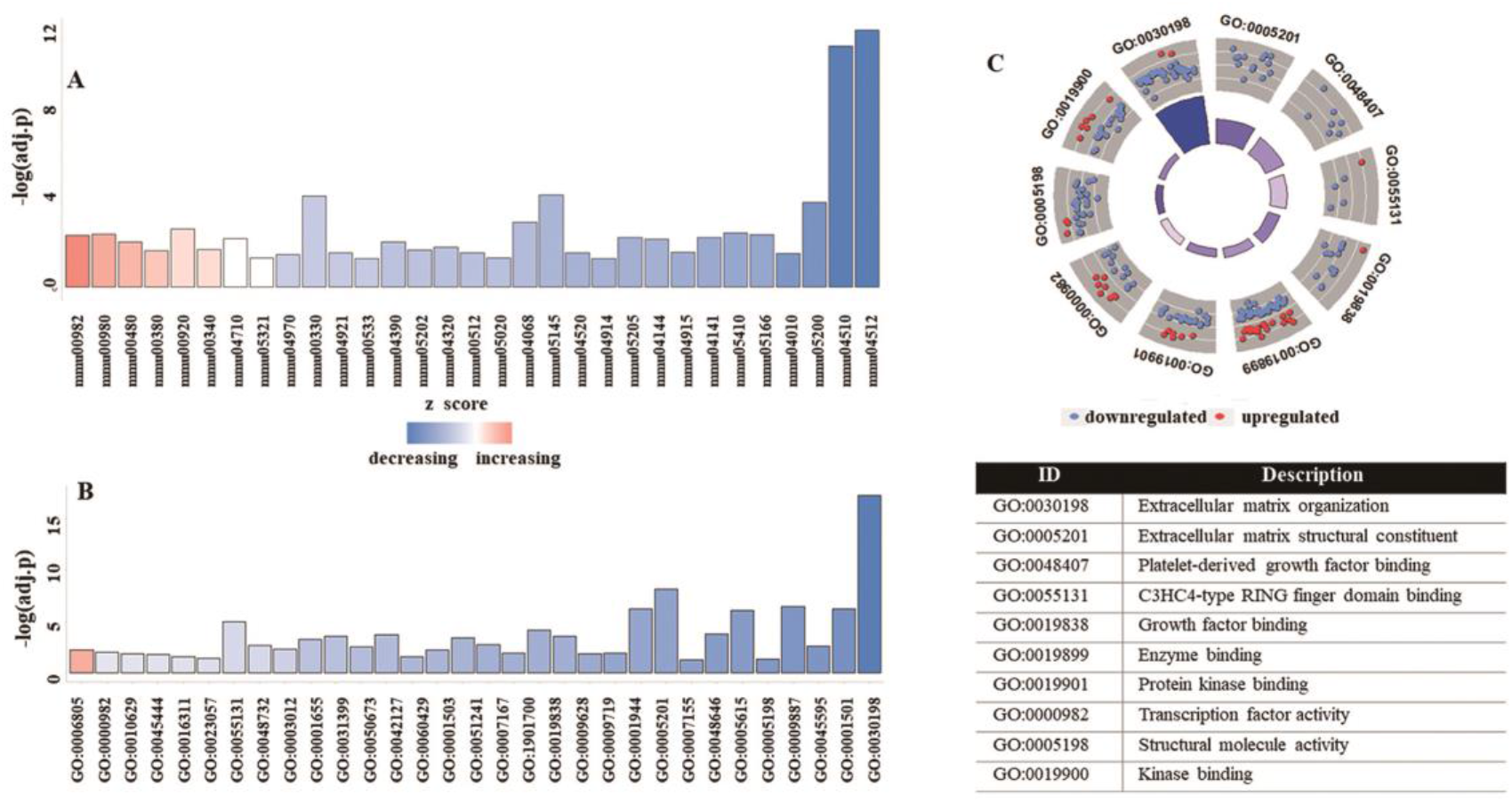
The representation of enrichment analysis results of common DEGs using GOplot. (A): The identified pathways (colors stand for corresponding z-score). (B): The identified functional annotation clusters (colors stand for corresponding z-score. (C): The circle picture of 10 most significant clusters where blue dots stand for down-regulated genes and red dots stand for up-regulated genes

**Fig. 5.**
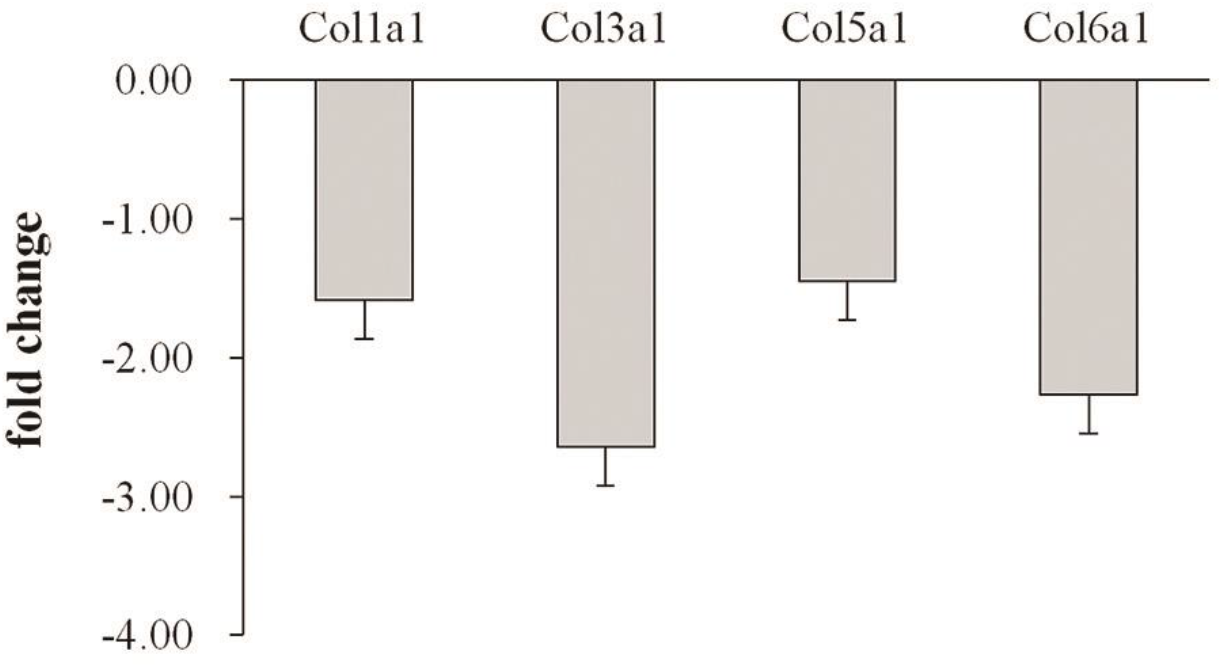
The fold changes of selected genes in skeletal muscles from mice exposed to normoxia (n=6) or 8% O_2_ for 2 hours (n=7)

### The potential relationship between metabolic changes and transcriptional changes evaluated by genome-based metabolic network reconstruction

According to the central dogma, the level of enzymes can be regulated by the level of its mRNA through translation. However, during the acute hypoxia duration, the level of mRNA is continuously changing. So, it is interesting whether such a change on mRNA can be translated into its enzyme and hence affects relevant metabolites during acute hypoxia. According to mouse metabolic flux network model iMM1415, we identified 49 differentially expressed metabolic genes and 176 metabolic reactions as listed in Table S5. Among these genes, genes encoding enzymes for amino acid metabolism were relatively enriched, which included Gcat, Glul, Gpt2, Dot1l, Bckdhb, Aldh2, Dpysl3, P4ha1, Aldh9a1, Gnmt and Gatm. Based on the assumption that the gene changes were proportionally translated into protein levels, we predicted the effects of metabolic cDEGs on metabolism which were visualized in Fig. S1. The predicted changes were sparsely distributed in metabolic subsystems, such as glutamine, tryptophan, cysteine and cytochrome p450 metabolism. The predicted change on glutamine metabolism was consistent with the increase in glutamic acid measured by LC-MS/MS, with increases in glutamine synthetase gene Glu1 and glutamic pyruvate transaminase gene Gpt2. However, a bit of predictions based on metabolic cDEGs such as Dpysl3, Gatm were not reflected by the metabolic measurement results. Compared with the hundreds of changes on gene expression, the changes on metabolites were quite moderate. Thus, the correlation between changes on mRNA levels and changes on relevant metabolites was not high. Thus, for hours of hypoxia, some changes on mRNA levels might not be reflected by protein levels timely as assumed. This means that we may need more mechanisms besides central dogma to interpret the results we observed in acute hypoxic exposure.

## Discussion

### Glutamine metabolism and lactate metabolism in skeletal muscles responding to hypoxic stress

Lactate metabolism was widely known as the most significant response to heavy exercises in skeletal muscles. A ~330% increase in blood lactate, which was recovered to the normal level in 2 hours, was observed after a 15-minute sedentary hypoxia along with significant decreases in blood PaO_2_, PaCO_2_ [11]. In contrast to rapid metabolic responses in blood, the lactate in plantaris muscles didn’t show a significant change either in 15 minutes or 2 hours of hypoxic exposure. It was known that skeletal muscles were the major source of increased blood lactate during high intensity exercise due to lactate accumulation in skeletal muscles [29]. However, the result of lactate in this study suggested that skeletal muscles wouldn’t be a major contributor of the increased blood lactate in a sedentary hypoxic exposure. This difference in skeletal muscles for lactate metabolism under the hypoxic stress induced by a bout exercise or a sedentary hypoxia was interesting. A sedentary hypoxia induced a ‘passive’ hypoxia in skeletal muscles which was delivered from the around tissue, e.g. from blood to skeletal muscles, while an exercise bout induced an ‘active’ hypoxia in skeletal muscles which was delivered to nearby tissues. Duarte et al. suggested brain as a potential source of increased blood lactate under hypoxia since a stable increase in the lactate release was observed in hippocampal slices in a hypoxic duration [30]. But this might not be the case in this study, since the level of blood lactate was recovered to the normal level when hypoxia was still last. Red blood cells were reported as one of the main lactate sources, which were sensitive to changes on blood oxygen level [31]. During the sedentary hypoxia, red blood cells sense the decrease of oxygen level much earlier than skeletal muscles, and hence might be the major source of blood lactate during a sedentary hypoxic exposure.

In this study, the increase in glutamic acid was the most significant metabolic change (22.05±4.04 v.s. 43.27±6.35) in skeletal muscles after 2 hours of hypoxia. Previous studies had identified a role of glutamine metabolism in hypoxia. Huang et al. found that glutamine played an important role in the growth of cancer cells which usually grew under hypoxic conditions [32]. The glutamine metabolic pathway was identified as the required pathway for tumor growth which rendered glutamine mainly to acetyl coenzyme A for lipogenesis [33]. Fuhrmann et al. reported that extended hypoxia in myocytes increased fatty acid oxidation via the glutamine-citrate-fatty acid axis [34]. Starnes et al. reported that glutamic acid and glutamine changed in heart during exercise inducing a temporary hypoxic condition [7]. In this study, we observed a rapid increase in glutamic acid in mouse skeletal muscles in a 2-hour hypoxic exposure. Thus, based on above studies, different from lactate metabolism, glutamine metabolism is widely involved in kinds of hypoxia-relevant events, such as sedentary hypoxia, hypoxia induced by a heavy exercise and hypoxia caused by the excessive growth in cancer. This fact suggests that glutamine metabolism might be a general responsive mechanism to hypoxic stress.

### Metabolites and gene expression levels in responses to acute sedantary hypoxia

Several studies had carried out to study the potential relationship between metabolomics and transcriptomics. Kelly et al. studied asthma by integrating metabolites, gene transcripts and found a role of dysregulation of lipid metabolism in asthma [35]. Feidantsis’s study indicated a close correlation between the expression of heat shock protein and the level of lipid peroxidation induced by a long-term thermal stress in mussels [36]. In this study, the genome-based metabolic network analysis predicted some measured metabolic changes such as glutamic acid, but also changes which couldn’t be reflected by measured metabolites. As an acute hypoxia, the expression levels of genes or proteins were perhaps changing continuously during the period. The rates of changes for genes or proteins could have different time scale and the translation of changes on genes into that of proteins would need time, thus it might not be practical to predict metabolites for the unstable state before we knew all dynamic and kinetic properties of genes, proteins and metabolites. Another possible reason could be that the traditional RNA-enzymatic protein-metabolism dogma, which was the assumption of the analysis method, may not be sufficient to interpret gene regulation mechanism under acute hypoxia. Other mechanisms such as epitranscriptome including methylation could be involved in gene expression regulation under acute hypoxic state. In this study, metabolite result indicated that 2 hours of hypoxia induced decreases in uridine and cytidine. Uridine was reported as a protective treatment for brain damage caused by hypoxic-ischemia [37]. A previous study showed that hypoxia could cause hundreds of RNA changes responding to hypoxia through APOBEC3A, a cytidine deaminase [38]. These studies suggested a potential role of uridine and cytidine in responses to hypoxia. The decreases in uridine and cytidine found in this study also suggested an involvement of uridine and cytidine in acute hypoxic responses, though the low levels of uridine and cytidine that we detected in skeletal muscles might affect the reliability of this point.

### Effects of tools on data alignment and differentially expressed gene identification

In this study, we tried different tools to analyze RNA-seq data with mouse reference genome mm9 or mm10. The selection of different reference genome indicated very slight effects on the results. The alignment to mm10 reported ~28 genes more than mm9. The gene counts and folds using either mm9 or mm10 were quite close, with an average ratio around 1.00±0.05. Thus, the selection of different versions of reference genome wouldn’t cause an essential difference on results.

We used TopHat2/Bowtie2 or Hisat2/StringTie to align and assemble the sequencing data. Hisat2 indicated a much faster alignment speed where the running time was decreased significantly. For an example, TopHat2 took about 220 minutes with 12 threads to align 30 Gb data to mm10, while it took only 15 minutes for Hisat2 with 12 threads to align the same input data to mm10. The alignment rate of Hisat2 was also slightly higher. The average of alignment rate by Hisat2 of our data was around 95%, while TopHat2 gave an alignment rate around 87%. Thus, Hisat2/StringTie had a better performance for the alignment. This conclusion was consistent with that of Sahraeian et al., who pointed out that Hisat2/StringTie were the fastest and most effective tools to align short reads [39].

To identify the effects of tool on the identification of DEGs, we tried Cufflinks, Deseq2 and AltAnalyze (limma-based). Cufflinks, Deseq2 and AltAnalyze identified 1057, 825 and 767 DEGs respectively. However, only 350 genes were common among them. The venn diagram in Fig. 4 showed the relationship of DEGs selected by different tools, suggesting that the selection of tools could have a big effect on the identification of DEGs. Fortunately, the folds of cDEGs calculated by different tools were quite comparable (Table S2). To coarsely evaluate these 3 sets of DEGs, we compared top 100 genes with most significant p or q values from 3 sets and top 100 genes with biggest fold changes from 3 sets. The results showed that DEGs identified by Cuffdiff had the least overlap with cDEGs as listed in Table 2. Another point that we noticed was that none of 10 DEGs with the biggest folds identified by Cuffdiff were recognized by DESeq2 or AltAnalyze, which might suggest a false discovery. This result also agreed with the conclusion of Sahraeian et al. who pointed out that outputs of pipelines involving Cufflinks/Cuffdiff had a less similarity to other pipeline combinations [39]. Selected by p value, DESeq2 seemed the most effective with 89 genes overlapped with cDEGs. AltAnalyze had a good performance at both the selection by p/q value and the selection by fold as shown in Table 2. Based above results, AltAnalyze or DESeq2 was better to identify differentially expressed genes rather than Cuffdiff. Though DEGs identified by different tools were different, we found their results of enrichment analyses such as pathway analysis were similar, suggesting that key differentially expressed genes were conserved among 3 sets. Thus, we used cDEGs for further analyses, to simplify the complexity of analyses and to focus on key information.

**Table 2.** The overlap among gene sets selected by different tools. Column 2 indicated the conservation rate of top 100 genes selected by p (AltAnalyze, DESeq2) or q values (Cuffdiff) compared to cDEGs. Column3 showed the conservation rate of top 100 genes selected by the absolute fold changes compared to cDEGs

| tool | by p/q value | by fold |
| --- | --- | --- |
| Cuffdiff | 38/100 | 16/100 |
| AltAnalyze | 73/100 | 76/100 |
| DESeq2 | 89/100 | 30/100 |

## Additional files

**Fig. S1.** The visualization of cDEGs and their changes in the genome-based mouse metabolic network

**Table S1.** The list of primers used for RT-PCR in this study

**Table S2.** The list of identified DEGs and their pathway analysis results

**Table S3 & S4.** The pathway and functional annotation clustering analysis result of cDEGs

**Table S5.** Mapped cDEGs and reactions in the genome-based mouse metabolic

## List of abbreviations

cDEG: conserved differentially expressed gene
DEG: differentially expressed gene

## Acknowledgements

The authors sincerely thank Drs. A.D. McCulloch, F.L Powell, D.C. Zielinski and Ms. Jennifer Stowe from University of California San Diego for their supports at the initial stage of this study. We also appreciate Biomedical Genetics and Metabolomics Laboratory, especially Dr. J.A. Gangoiti from University of California San Diego who helped us design and conduct the metabolite measurements.

## Author contribution

Z. Gan conceived and designed this study. Z. Gan, J. Deng, X Pang and F. Xue contributed to the animal and molecular experiments. Z. Gan, G. Huang and X. Wang conducted bioinformatics analyses. Z. Gan drafted the manuscript. All authors approved the manuscript.

## Ethics approval and consent to participate

All animal procedures were approved by Animal Care and Use Committee of Wenzhou Medical University.

## References

1. Gonzalez FJ, Xie C, Jiang C. The role of hypoxia-inducible factors in metabolic diseases. Nat Rev Endocrinol. 2018; 15(1):21–32.

2. Chaillou T. Skeletal Muscle Fiber Type in Hypoxia: Adaptation to High-Altitude Exposure and Under Conditions of Pathological Hypoxia. Front Physiol. 2018; 9:1450.

3. Nanduri J, Semenza GL, Prabhakar NR. Epigenetic changes by DNA methylation in chronic and intermittent hypoxia. Am J Physiol Lung Cell Mol Physiol. 2017; 313(6):L1096–L1100.

4. Burtscher M, Gatterer H, Burtscher J, Mairbaurl H. Extreme Terrestrial Environments: Life in Thermal Stress and Hypoxia. A Narrative Review. Front Physiol. 2018; 9:572.

5. Morales-Alamo D, Guerra B, Santana A, Martin-Rincon M, Gelabert-Rebato M, Dorado C et al. Skeletal Muscle Pyruvate Dehydrogenase Phosphorylation and Lactate Accumulation During Sprint Exercise in Normoxia and Severe Acute Hypoxia: Effects of Antioxidants. Front Physiol. 2018; 9:188.

6. Hoene M, Li J, Li Y, Runge H, Zhao X, Haring HU et al. Muscle and liver-specific alterations in lipid and acylcarnitine metabolism after a single bout of exercise in mice. Sci Rep. 2016; 6:22218.

7. Starnes JW, Parry TL, O’Neal SK, Bain JR, Muehlbauer MJ, Honcoop A et al. Exercise-Induced Alterations in Skeletal Muscle, Heart, Liver, and Serum Metabolome Identified by Non-Targeted Metabolomics Analysis. Metabolites. 2017; 7(3).

8. Kim SH, Koh JH, Higashida K, Jung SR, Holloszy JO, Han DH. PGC-1alpha mediates a rapid, exercise-induced downregulation of glycogenolysis in rat skeletal muscle. J Physiol. 2015; 593(3):635–643.

9. Azad P, Zhou D, Russo E, Haddad GG. Distinct mechanisms underlying tolerance to intermittent and constant hypoxia in Drosophila melanogaster. PLoS One. 2009; 4(4):e5371.

10. Wollen EJ, Sejersted Y, Wright MS, Madetko-Talowska A, Bik-Multanowski M, Kwinta P et al. Transcriptome profiling of the newborn mouse brain after hypoxia-reoxygenation: hyperoxic reoxygenation induces inflammatory and energy failure responsive genes. Pediatr Res. 2014; 75(4):517–526.

11. Gan Z, Powell FL, Zambon AC, Buchholz KS, Fu Z, Ocorr K et al. Transcriptomic analysis identifies a role of PI3K-Akt signalling in the responses of skeletal muscle to acute hypoxia in vivo. J Physiol. 2017; 595(17):5797–5813.

12. Sukhanova IA, Sebentsova EA, Khukhareva DD, Manchenko DM, Glazova NY, Vishnyakova PA et al. Gender-dependent changes in physical development, BDNF content and GSH redox system in a model of acute neonatal hypoxia in rats. Behav Brain Res. 2018; 350:87–98.

13. Qi D, Xia M, Chao Y, Zhao Y, Wu R. Identification, molecular evolution of toll-like receptors in a Tibetan schizothoracine fish (Gymnocypris eckloni) and their expression profiles in response to acute hypoxia. Fish Shellfish Immunol. 2017; 68:102–113.

14. Bartoszewski R, Serocki M, Janaszak-Jasiecka A, Bartoszewska S, Kochan-Jamrozy K, Piotrowski A et al. miR-200b downregulates Kruppel Like Factor 2 (KLF2) during acute hypoxia in human endothelial cells. Eur J Cell Biol. 2017; 96(8):758–766.

15. Burkholder TJ, Fingado B, Baron S, Lieber RL. Relationship between muscle fiber types and sizes and muscle architectural properties in the mouse hindlimb. J Morphol. 1994; 221(2):177–190.

16. Gan Z, Fu Z, Stowe JC, Powell FL, McCulloch AD. A Protocol to Collect Specific Mouse Skeletal Muscles for Metabolomics Studies. Methods Mol Biol. 2016; 1375:169–179.

17. Gertsman I, Gangoiti JA, Barshop BA. Validation of a dual LC-HRMS platform for clinical metabolic diagnosis in serum, bridging quantitative analysis and untargeted metabolomics. Metabolomics. 2014; 10(2):312–323.

18. Pertea M, Kim D, Pertea GM, Leek JT, Salzberg SL. Transcript-level expression analysis of RNA-seq experiments with HISAT, StringTie and Ballgown. Nat Protoc. 2016; 11(9):1650–1667.

19. Emig D, Salomonis N, Baumbach J, Lengauer T, Conklin BR, Albrecht M. AltAnalyze and DomainGraph: analyzing and visualizing exon expression data. Nucleic Acids Res. 2010; 38(Web Server issue):W755–762.

20. Love MI, Huber W, Anders S. Moderated estimation of fold change and dispersion for RNA-seq data with DESeq2. Genome Biol. 2014; 15(12):550.

21. Trapnell C, Hendrickson DG, Sauvageau M, Goff L, Rinn JL, Pachter L. Differential analysis of gene regulation at transcript resolution with RNA-seq. Nat Biotechnol. 2013; 31(1):46–53.

22. Chen J, Bardes EE, Aronow BJ, Jegga AG. ToppGene Suite for gene list enrichment analysis and candidate gene prioritization. Nucleic Acids Res. 2009; 37(Web Server issue):W305–311.

23. Walter W, Sanchez-Cabo F, Ricote M. GOplot: an R package for visually combining expression data with functional analysis. Bioinformatics. 2015; 31(17):2912–2914.

24. King ZA, Drager A, Ebrahim A, Sonnenschein N, Lewis NE, Palsson BO. Escher: A Web Application for Building, Sharing, and Embedding Data-Rich Visualizations of Biological Pathways. PLoS Comput Biol. 2015; 11(8):e1004321.

25. Sigurdsson MI, Jamshidi N, Steingrimsson E, Thiele I, Palsson BO. A detailed genome-wide reconstruction of mouse metabolism based on human Recon 1. BMC Syst Biol. 2010; 4:140.

26. Mattei G, Zielinski DC, Gan Z, Ramazzotti M, Palsson BO. Functional pathways for metabolic network-based data analysis: the MetPath algorithm. bioRxiv. 2017.

27. Vianna CR, Huntgeburth M, Coppari R, Choi CS, Lin J, Krauss S et al. Hypomorphic mutation of PGC-1beta causes mitochondrial dysfunction and liver insulin resistance. Cell Metab. 2006; 4(6):453–464.

28. Wang Y, Eddy JA, Price ND. Reconstruction of genome-scale metabolic models for 126 human tissues using mCADRE. BMC Syst Biol. 2012; 6:153.

29. Nugent WH, Song BK, Pittman RN, Golub AS. Simultaneous sampling of tissue oxygenation and oxygen consumption in skeletal muscle. Microvasc Res. 2016; 105:15–22.

30. Duarte JM, Cunha RA, Carvalho RA. Adenosine A(1) receptors control the metabolic recovery after hypoxia in rat hippocampal slices. J Neurochem. 2016; 136(5):947–957.

31. Seheult J, Fitzpatrick G, Boran G. Lactic acidosis: an update. Clin Chem Lab Med. 2017; 55(3):322–333.

32. Huang W, Choi W, Chen Y, Zhang Q, Deng H, He W et al. A proposed role for glutamine in cancer cell growth through acid resistance. Cell Res. 2013; 23(5):724–727.

33. Wang Y, Bai C, Ruan Y, Liu M, Chu Q, Qiu L et al. Coordinative metabolism of glutamine carbon and nitrogen in proliferating cancer cells under hypoxia. Nat Commun. 2019; 10(1):201.

34. Fuhrmann DC, Olesch C, Kurrle N, Schnutgen F, Zukunft S, Fleming I et al. Chronic Hypoxia Enhances beta-Oxidation-Dependent Electron Transport via Electron Transferring Flavoproteins. Cells. 2019; 8(2).

35. Kelly RS, Chawes BL, Blighe K, Virkud YV, Croteau-Chonka DC, McGeachie MJ et al. An Integrative Transcriptomic and Metabolomic Study of Lung Function in Children With Asthma. Chest. 2018; 154(2):335–348.

36. Feidantsis K, Giantsis IA, Vratsistas A, Makri S, Pappa AZ, Drosopoulou E et al. Correlation between intermediary metabolism, Hsp gene expression, and oxidative stress-related proteins in long-term thermal-stressed Mytilus galloprovincialis. Am J Physiol Regul Integr Comp Physiol. 2020; 319(3):R264–R281.

37. Goren B, Cakir A, Ocalan B, Serter Kocoglu S, Alkan T, Cansev M et al. Long-term cognitive effects of uridine treatment in a neonatal rat model of hypoxic-ischemic encephalopathy. Brain Res. 2017; 1659:81–87.

38. Sharma S, Patnaik SK, Taggart RT, Kannisto ED, Enriquez SM, Gollnick P et al. APOBEC3A cytidine deaminase induces RNA editing in monocytes and macrophages. Nat Commun. 2015; 6:6881.

39. Sahraeian SME, Mohiyuddin M, Sebra R, Tilgner H, Afshar PT, Au KF et al. Gaining comprehensive biological insight into the transcriptome by performing a broad-spectrum RNA-seq analysis. Nat Commun. 2017; 8(1):59.

