## Supplementary figures and images for "An exploration of metabolite and gene responses in mouse skeletal muscles responding to acute sedentary hypoxia"

### Fig. S1

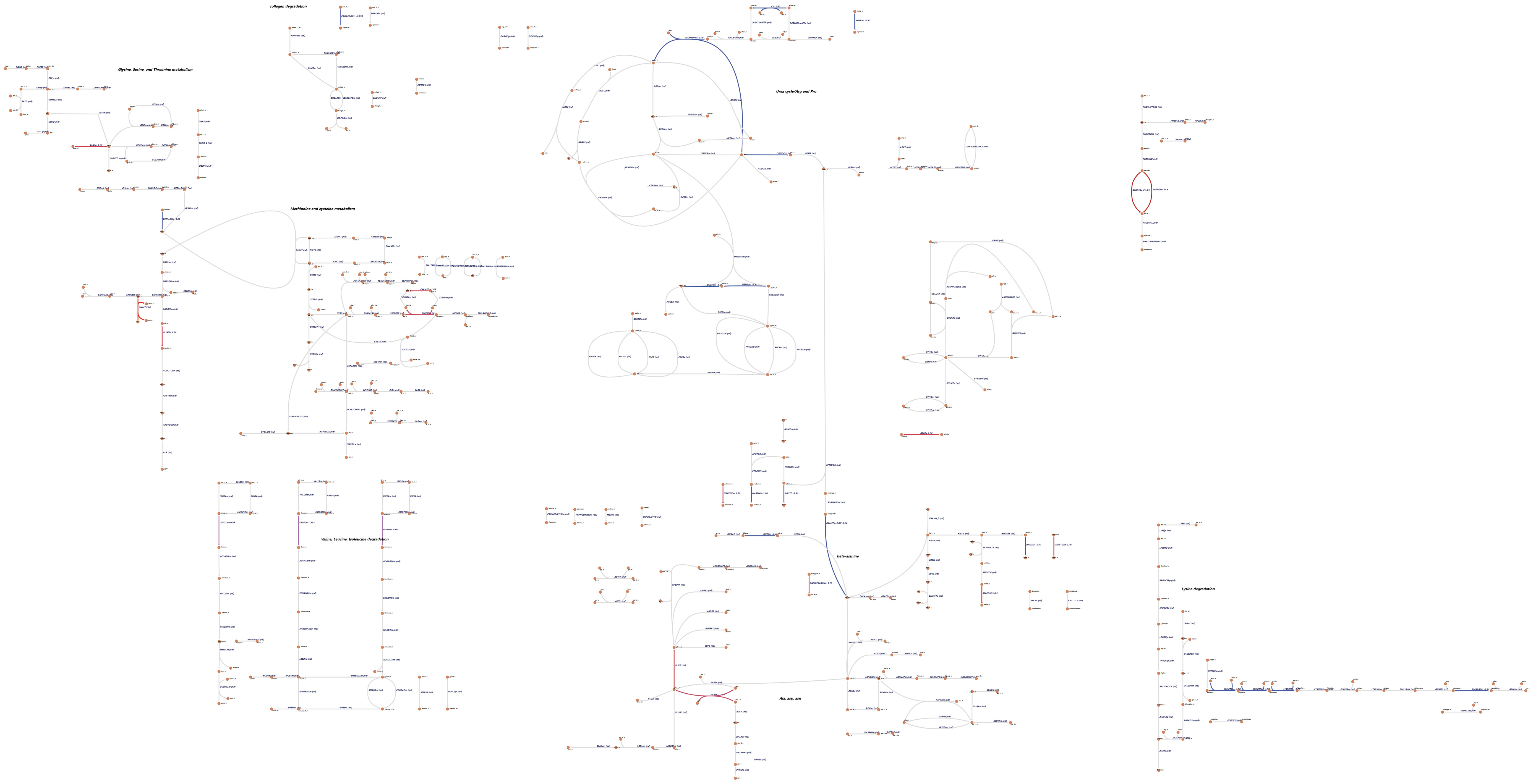
