## Supplementary material for "An exploration of metabolite and gene responses in mouse skeletal muscles responding to acute sedentary hypoxia": Table S1

| The list of primers used for RT-PCR in this study | | |
|---|---|---|
| **Gene** | **Forward** | **Reverse** |
| 8s | GTAACCCGTTGAACCCCATT | CCATCCAATCGGTAGTAGCG |
| Col1a1 | GCTCCTCTTAGGGGCCACT | CCACGTCTCACCATTGGGG |
| Col3a1 | CTGTAACATGGAAACTGGGGAAA | CCATAGCTGAACTGAAAACCAC |
| Col5a1 | TGAGTCTGGTTTTCCCGAGGA | GCCCTGCTCATTGTAAATGGAGA |
| Col6a1 | CTGCTGCTACAAGCCTGCT | CCCCATAAGGTTTCAGCCTCA |
