## Supplementary material for "An exploration of metabolite and gene responses in mouse skeletal muscles responding to acute sedentary hypoxia": Table S2

| pathway analysis result of DEGs identified by different tools | | | |
|---|---|---|---|
| pathway | #DEGs | #total targets | p |
| **tool: AltAnalyze** | | | |
| ECM-receptor interaction | 21 | 88 | 1.17E-11 |
| Focal adhesion | 30 | 207 | 3.14E-10 |
| Amoebiasis | 20 | 119 | 2.55E-08 |
| Protein digestion and absorption | 16 | 89 | 2.53E-07 |
| Arginine and proline metabolism | 11 | 59 | 1.37E-05 |
| Glutathione metabolism | 10 | 55 | 4.26E-05 |
| PI3K-Akt signaling pathway | 29 | 351 | 9.41E-05 |
| Metabolism of xenobiotics by cytochrome P450 | 10 | 64 | 1.62E-04 |
| Platelet activation | 15 | 131 | 1.63E-04 |
| Drug metabolism - cytochrome P450 | 10 | 66 | 2.10E-04 |
| Chemical carcinogenesis | 12 | 92 | 2.17E-04 |
| Toxoplasmosis | 13 | 113 | 4.23E-04 |
| Lysine degradation | 8 | 51 | 7.13E-04 |
| Circadian rhythm | 6 | 31 | 1.07E-03 |
| Small cell lung cancer | 10 | 85 | 1.63E-03 |
| Pathways in cancer | 24 | 323 | 1.71E-03 |
| Hippo signaling pathway | 13 | 154 | 6.94E-03 |
| Sulfur metabolism | 3 | 11 | 7.65E-03 |
| Tryptophan metabolism | 6 | 46 | 8.38E-03 |
| Endocytosis | 17 | 231 | 8.61E-03 |
| Prion diseases | 5 | 35 | 1.08E-02 |
| AMPK signaling pathway | 11 | 129 | 1.17E-02 |
| Histidine metabolism | 4 | 24 | 1.29E-02 |
| Pancreatic cancer | 7 | 66 | 1.38E-02 |
| TGF-beta signaling pathway | 8 | 82 | 1.41E-02 |
| Protein processing in endoplasmic reticulum | 13 | 169 | 1.45E-02 |
| Proteoglycans in cancer | 16 | 226 | 1.50E-02 |
| FoxO signaling pathway | 11 | 135 | 1.60E-02 |
| Hypertrophic cardiomyopathy (HCM) | 8 | 84 | 1.62E-02 |
| Pyruvate metabolism | 5 | 39 | 1.69E-02 |
| Dilated cardiomyopathy | 8 | 89 | 2.22E-02 |
| HTLV-I infection | 18 | 277 | 2.27E-02 |
| Legionellosis | 6 | 58 | 2.46E-02 |
| Nitrogen metabolism | 3 | 17 | 2.65E-02 |
| beta-Alanine metabolism | 4 | 32 | 3.44E-02 |
| Colorectal cancer | 6 | 64 | 3.76E-02 |
| Fatty acid degradation | 5 | 48 | 3.79E-02 |
| MAPK signaling pathway | 16 | 253 | 3.80E-02 |
| Glycolysis / Gluconeogenesis | 6 | 65 | 4.01E-02 |
| Hedgehog signaling pathway | 5 | 49 | 4.09E-02 |
| Transcriptional misregulation in cancer | 12 | 178 | 4.50E-02 |
| Metabolic pathways | 60 | 1254 | 4.72E-02 |
| **tool: cuffdiff** | | | |
| Focal adhesion | 46 | 207 | 3.43E-14 |
| ECM-receptor interaction | 28 | 88 | 3.74E-13 |
| Protein digestion and absorption | 24 | 89 | 8.74E-10 |
| Amoebiasis | 26 | 119 | 2.22E-08 |
| Phagosome | 32 | 174 | 4.40E-08 |

| | | | |
|---|---|---|---|
| Toxoplasmosis | 23 | 113 | 5.65E-07 |
| PI3K-Akt signaling pathway | 47 | 351 | 8.86E-07 |
| Circadian rhythm | 11 | 31 | 1.78E-06 |
| Platelet activation | 24 | 131 | 2.37E-06 |
| Staphylococcus aureus infection | 13 | 51 | 1.33E-05 |
| Proteoglycans in cancer | 31 | 226 | 4.30E-05 |
| Osteoclast differentiation | 21 | 126 | 4.50E-05 |
| Tuberculosis | 26 | 176 | 5.06E-05 |
| Cytokine-cytokine receptor interaction | 34 | 264 | 6.83E-05 |
| Hypertrophic cardiomyopathy (HCM) | 16 | 84 | 7.06E-05 |
| Legionellosis | 12 | 58 | 2.52E-04 |
| Inflammatory bowel disease (IBD) | 12 | 59 | 2.98E-04 |
| Fc gamma R-mediated phagocytosis | 15 | 88 | 4.24E-04 |
| Dilated cardiomyopathy | 15 | 89 | 4.80E-04 |
| Mineral absorption | 10 | 46 | 5.40E-04 |
| Glutathione metabolism | 11 | 55 | 6.15E-04 |
| FoxO signaling pathway | 19 | 135 | 9.43E-04 |
| Hematopoietic cell lineage | 14 | 86 | 1.05E-03 |
| Prion diseases | 8 | 35 | 1.37E-03 |
| HTLV-I infection | 31 | 277 | 1.62E-03 |
| Leukocyte transendothelial migration | 17 | 121 | 1.75E-03 |
| Hippo signaling pathway | 20 | 154 | 1.92E-03 |
| Leishmaniasis | 11 | 65 | 2.57E-03 |
| Pathways in cancer | 34 | 323 | 2.77E-03 |
| Malaria | 9 | 48 | 3.04E-03 |
| Salivary secretion | 12 | 77 | 3.41E-03 |
| Gap junction | 13 | 87 | 3.44E-03 |
| p53 signaling pathway | 11 | 68 | 3.71E-03 |
| Arginine and proline metabolism | 10 | 59 | 3.94E-03 |
| Transcriptional misregulation in cancer | 21 | 178 | 4.81E-03 |
| Adipocytokine signaling pathway | 11 | 71 | 5.20E-03 |
| Antigen processing and presentation | 12 | 81 | 5.21E-03 |
| Regulation of actin cytoskeleton | 24 | 217 | 6.10E-03 |
| Measles | 17 | 137 | 6.48E-03 |
| Endocytosis | 25 | 231 | 6.83E-03 |
| Small cell lung cancer | 12 | 85 | 7.68E-03 |
| Adrenergic signaling in cardiomyocytes | 18 | 151 | 7.83E-03 |
| AMPK signaling pathway | 16 | 129 | 8.16E-03 |
| Complement and coagulation cascades | 11 | 77 | 9.61E-03 |
| Viral myocarditis | 11 | 78 | 1.06E-02 |
| Oxytocin signaling pathway | 18 | 158 | 1.23E-02 |
| cGMP-PKG signaling pathway | 19 | 170 | 1.25E-02 |
| Influenza A | 19 | 170 | 1.25E-02 |
| Chagas disease (American trypanosomiasis) | 13 | 103 | 1.42E-02 |
| Bile secretion | 10 | 72 | 1.60E-02 |
| Pertussis | 10 | 74 | 1.91E-02 |
| Metabolism of xenobiotics by cytochrome P450 | 9 | 64 | 2.01E-02 |
| Jak-STAT signaling pathway | 17 | 155 | 2.07E-02 |
| Estrogen signaling pathway | 12 | 98 | 2.24E-02 |
| Drug metabolism - cytochrome P450 | 9 | 66 | 2.42E-02 |
| HIF-1 signaling pathway | 13 | 111 | 2.50E-02 |
| Prostate cancer | 11 | 89 | 2.64E-02 |
| Sulfur metabolism | 3 | 11 | 2.94E-02 |
| Thyroid hormone signaling pathway | 13 | 118 | 3.87E-02 |

| | | | |
|---|---|---|---|
| Glycine serine and threonine metabolism | 6 | 40 | 4.05E-02 |
| Dorso-ventral axis formation | 4 | 22 | 4.88E-02 |
| **tool: DESeq2** | | | |
| Focal adhesion | 37 | 207 | 1.66E-13 |
| ECM-receptor interaction | 24 | 88 | 2.67E-13 |
| Amoebiasis | 21 | 119 | 4.53E-08 |
| Protein digestion and absorption | 15 | 89 | 7.14E-06 |
| Arginine and proline metabolism | 12 | 59 | 8.15E-06 |
| PI3K-Akt signaling pathway | 34 | 351 | 1.25E-05 |
| Platelet activation | 18 | 131 | 1.71E-05 |
| Toxoplasmosis | 16 | 113 | 3.45E-05 |
| Small cell lung cancer | 13 | 85 | 8.38E-05 |
| Pathways in cancer | 30 | 323 | 9.06E-05 |
| Circadian rhythm | 7 | 31 | 3.33E-04 |
| Proteoglycans in cancer | 22 | 226 | 4.18E-04 |
| Hypertrophic cardiomyopathy (HCM) | 11 | 84 | 1.13E-03 |
| Dilated cardiomyopathy | 11 | 89 | 1.83E-03 |
| Glutathione metabolism | 8 | 55 | 2.71E-03 |
| Protein processing in endoplasmic reticulum | 16 | 169 | 3.32E-03 |
| Legionellosis | 8 | 58 | 3.81E-03 |
| Oxytocin signaling pathway | 15 | 158 | 4.29E-03 |
| HTLV-I infection | 22 | 277 | 5.66E-03 |
| Thyroid hormone signaling pathway | 12 | 118 | 5.94E-03 |
| Cytokine-cytokine receptor interaction | 21 | 264 | 6.68E-03 |
| Endocytosis | 19 | 231 | 6.83E-03 |
| Lysine degradation | 7 | 51 | 6.85E-03 |
| Glycolysis / Gluconeogenesis | 8 | 65 | 7.67E-03 |
| Sulfur metabolism | 3 | 11 | 1.10E-02 |
| Estrogen signaling pathway | 10 | 98 | 1.13E-02 |
| Transcriptional misregulation in cancer | 15 | 178 | 1.25E-02 |
| beta-Alanine metabolism | 5 | 32 | 1.25E-02 |
| Adipocytokine signaling pathway | 8 | 71 | 1.29E-02 |
| Inflammatory bowel disease (IBD) | 7 | 59 | 1.49E-02 |
| FoxO signaling pathway | 12 | 135 | 1.65E-02 |
| Measles | 12 | 137 | 1.83E-02 |
| Hippo signaling pathway | 13 | 154 | 1.91E-02 |
| Histidine metabolism | 4 | 24 | 2.00E-02 |
| Biosynthesis of amino acids | 8 | 78 | 2.17E-02 |
| Metabolism of xenobiotics by cytochrome P450 | 7 | 64 | 2.26E-02 |
| Drug metabolism - cytochrome P450 | 7 | 66 | 2.62E-02 |
| Pancreatic cancer | 7 | 66 | 2.62E-02 |
| AMPK signaling pathway | 11 | 129 | 2.79E-02 |
| Pyruvate metabolism | 5 | 39 | 2.79E-02 |
| TGF-beta signaling pathway | 8 | 82 | 2.84E-02 |
| Rheumatoid arthritis | 8 | 82 | 2.84E-02 |
| Ascorbate and aldarate metabolism | 4 | 27 | 2.98E-02 |
| Glycine serine and threonine metabolism | 5 | 40 | 3.07E-02 |
| Hematopoietic cell lineage | 8 | 86 | 3.64E-02 |
| Bile secretion | 7 | 72 | 3.97E-02 |
| Leukocyte transendothelial migration | 10 | 121 | 4.21E-02 |
| Arrhythmogenic right ventricular cardiomyopathy | 7 | 74 | 4.51E-02 |
