## Supplementary material for "An exploration of metabolite and gene responses in mouse skeletal muscles responding to acute sedentary hypoxia": Table S3

| The pathway analysis result of cDEGs | | | | |
|---|---|---|---|---|
| Pathway ID | Pathway name | #cDEGs | #totaltargets | p |
| mmu04512 | ECM-receptor interaction | 18 | 88 | 3.49E-13 |
| mmu04510 | Focal adhesion | 25 | 207 | 2.09E-12 |
| mmu05146 | Amoebiasis | 17 | 119 | 6.48E-10 |
| mmu04974 | Protein digestion and absorption | 13 | 89 | 5.60E-08 |
| mmu04151 | PI3K-Akt signaling pathway | 23 | 351 | 2.10E-06 |
| mmu05145 | Toxoplasmosis | 11 | 113 | 3.40E-05 |
| mmu00330 | Arginine and proline metabolism | 8 | 59 | 3.86E-05 |
| mmu05200 | Pathways in cancer | 19 | 323 | 7.72E-05 |
| mmu04611 | Platelet activation | 11 | 131 | 1.32E-04 |
| mmu05222 | Small cell lung cancer | 8 | 85 | 5.17E-04 |
| mmu04068 | FoxO signaling pathway | 10 | 135 | 7.29E-04 |
| mmu00920 | Sulfur metabolism | 3 | 11 | 1.50E-03 |
| mmu05134 | Legionellosis | 6 | 58 | 1.60E-03 |
| mmu05410 | Hypertrophic cardiomyopathy (HCM) | 7 | 84 | 2.36E-03 |
| mmu00980 | Metabolism of xenobiotics by cytochrome P450 | 6 | 64 | 2.66E-03 |
| mmu05166 | HTLV-I infection | 14 | 277 | 2.98E-03 |
| mmu00982 | Drug metabolism - cytochrome P450 | 6 | 66 | 3.11E-03 |
| mmu05414 | Dilated cardiomyopathy | 7 | 89 | 3.27E-03 |
| mmu04141 | Protein processing in endoplasmic reticulum | 10 | 169 | 3.91E-03 |
| mmu05205 | Proteoglycans in cancer | 12 | 226 | 3.95E-03 |
| mmu04710 | Circadian rhythm | 4 | 31 | 4.42E-03 |
| mmu04144 | Endocytosis | 12 | 231 | 4.71E-03 |
| mmu04390 | Hippo signaling pathway | 9 | 154 | 6.61E-03 |
| mmu00480 | Glutathione metabolism | 5 | 55 | 6.85E-03 |
| mmu04320 | Dorso-ventral axis formation | 3 | 22 | 1.17E-02 |
| mmu05210 | Colorectal cancer | 5 | 64 | 1.28E-02 |
| mmu05215 | Prostate cancer | 6 | 89 | 1.32E-02 |
| mmu00340 | Histidine metabolism | 3 | 24 | 1.49E-02 |
| mmu05204 | Chemical carcinogenesis | 6 | 92 | 1.53E-02 |
| mmu05202 | Transcriptional misregulation in cancer | 9 | 178 | 1.62E-02 |
| mmu00380 | Tryptophan metabolism | 4 | 46 | 1.78E-02 |
| mmu04122 | Sulfur relay system | 2 | 10 | 1.92E-02 |
| mmu04915 | Estrogen signaling pathway | 6 | 98 | 2.04E-02 |

| | | | | |
|---|---|---|---|---|
| mmu00071 | Fatty acid degradation | 4 | 48 | 2.05E-02 |
| mmu04921 | Oxytocin signaling pathway | 8 | 158 | 2.26E-02 |
| mmu00512 | Mucin type O-Glycan biosynthesis | 3 | 28 | 2.27E-02 |
| mmu04520 | Adherens junction | 5 | 74 | 2.28E-02 |
| mmu04010 | MAPK signaling pathway | 11 | 253 | 2.30E-02 |
| mmu00310 | Lysine degradation | 4 | 51 | 2.50E-02 |
| mmu04970 | Salivary secretion | 5 | 77 | 2.66E-02 |
| mmu00410 | beta-Alanine metabolism | 3 | 32 | 3.22E-02 |
| mmu05164 | Influenza A | 8 | 170 | 3.31E-02 |
| mmu04350 | TGF-beta signaling pathway | 5 | 82 | 3.36E-02 |
| mmu05321 | Inflammatory bowel disease (IBD) | 4 | 59 | 3.99E-02 |
| mmu05020 | Prion diseases | 3 | 35 | 4.06E-02 |
| mmu00533 | Glycosaminoglycan biosynthesis - keratan sulfate | 2 | 15 | 4.16E-02 |
| mmu04914 | Progesterone-mediated oocyte maturation | 5 | 87 | 4.18E-02 |
| mmu05212 | Pancreatic cancer | 4 | 66 | 5.63E-02 |
| mmu00260 | Glycine serine and threonine metabolism | 3 | 40 | 5.66E-02 |
| mmu00270 | Cysteine and methionine metabolism | 3 | 40 | 5.66E-02 |
| mmu04115 | p53 signaling pathway | 4 | 68 | 6.16E-02 |
| mmu04152 | AMPK signaling pathway | 6 | 129 | 6.38E-02 |
| mmu04060 | Cytokine-cytokine receptor interaction | 10 | 264 | 6.44E-02 |
| mmu05218 | Melanoma | 4 | 71 | 6.99E-02 |
| mmu04976 | Bile secretion | 4 | 72 | 7.28E-02 |
| mmu05220 | Chronic myeloid leukemia | 4 | 73 | 7.58E-02 |
| mmu05162 | Measles | 6 | 137 | 8.02E-02 |
| mmu05144 | Malaria | 3 | 48 | 8.75E-02 |
| mmu05416 | Viral myocarditis | 4 | 78 | 9.15E-02 |
| mmu03320 | PPAR signaling pathway | 4 | 81 | 1.02E-01 |
| mmu04612 | Antigen processing and presentation | 4 | 81 | 1.02E-01 |
| mmu01100 | Metabolic pathways | 34 | 1254 | 1.04E-01 |
| mmu00280 | Valine leucine and isoleucine degradation | 3 | 52 | 1.05E-01 |
| mmu04261 | Adrenergic signaling in cardiomyocytes | 6 | 151 | 1.14E-01 |
| mmu00053 | Ascorbate and aldarate metabolism | 2 | 27 | 1.17E-01 |
| mmu04640 | Hematopoietic cell lineage | 4 | 86 | 1.20E-01 |
| mmu04540 | Gap junction | 4 | 87 | 1.23E-01 |
| mmu00232 | Caffeine metabolism | 1 | 6 | 1.25E-01 |

| | | | | |
|---|---|---|---|---|
| mmu05216 | Thyroid cancer | 2 | 29 | 1.32E-01 |
| mmu04110 | Cell cycle | 5 | 126 | 1.43E-01 |
| mmu00760 | Nicotinate and nicotinamide metabolism | 2 | 32 | 1.55E-01 |
| mmu03040 | Spliceosome | 5 | 132 | 1.64E-01 |
| mmu04022 | cGMP-PKG signaling pathway | 6 | 170 | 1.70E-01 |
| mmu00010 | Glycolysis / Gluconeogenesis | 3 | 65 | 1.70E-01 |
| mmu05140 | Leishmaniasis | 3 | 65 | 1.70E-01 |
| mmu05214 | Glioma | 3 | 65 | 1.70E-01 |
| mmu05143 | African trypanosomiasis | 2 | 34 | 1.70E-01 |
| mmu05211 | Renal cell carcinoma | 3 | 67 | 1.81E-01 |
| mmu05169 | Epstein-Barr virus infection | 7 | 212 | 1.84E-01 |
| mmu00040 | Pentose and glucuronate interconversions | 2 | 36 | 1.86E-01 |
| mmu05142 | Chagas disease (American trypanosomiasis) | 4 | 103 | 1.89E-01 |
| mmu04015 | Rap1 signaling pathway | 7 | 214 | 1.90E-01 |
| mmu00250 | Alanine aspartate and glutamate metabolism | 2 | 37 | 1.94E-01 |
| mmu00072 | Synthesis and degradation of ketone bodies | 1 | 10 | 1.99E-01 |
| mmu04918 | Thyroid hormone synthesis | 3 | 71 | 2.04E-01 |
| mmu04920 | Adipocytokine signaling pathway | 3 | 71 | 2.04E-01 |
| mmu00620 | Pyruvate metabolism | 2 | 39 | 2.10E-01 |
| mmu05161 | Hepatitis B | 5 | 145 | 2.13E-01 |
| mmu04971 | Gastric acid secretion | 3 | 74 | 2.21E-01 |
| mmu04114 | Oocyte meiosis | 4 | 111 | 2.26E-01 |
| mmu05100 | Bacterial invasion of epithelial cells | 3 | 77 | 2.38E-01 |
| mmu01230 | Biosynthesis of amino acids | 3 | 78 | 2.44E-01 |
| mmu00061 | Fatty acid biosynthesis | 1 | 13 | 2.51E-01 |
| mmu00790 | Folate biosynthesis | 1 | 13 | 2.51E-01 |
| mmu04919 | Thyroid hormone signaling pathway | 4 | 118 | 2.59E-01 |
| mmu04146 | Peroxisome | 3 | 81 | 2.62E-01 |
| mmu00603 | Glycosphingolipid biosynthesis - globo series | 1 | 15 | 2.83E-01 |
| mmu00604 | Glycosphingolipid biosynthesis - ganglio series | 1 | 15 | 2.83E-01 |
| mmu04340 | Hedgehog signaling pathway | 2 | 49 | 2.92E-01 |
| mmu04380 | Osteoclast differentiation | 4 | 126 | 2.99E-01 |
| mmu00120 | Primary bile acid biosynthesis | 1 | 16 | 2.99E-01 |
| mmu04270 | Vascular smooth muscle contraction | 4 | 128 | 3.09E-01 |
| mmu00450 | Selenocompound metabolism | 1 | 17 | 3.14E-01 |

| | | | | |
|---|---|---|---|---|
| mmu00910 | Nitrogen metabolism | 1 | 17 | 3.14E-01 |
| mmu00983 | Drug metabolism - other enzymes | 2 | 52 | 3.16E-01 |
| mmu00511 | Other glycan degradation | 1 | 18 | 3.29E-01 |
| mmu00561 | Glycerolipid metabolism | 2 | 55 | 3.41E-01 |
| mmu00230 | Purine metabolism | 5 | 176 | 3.43E-01 |
| mmu01210 | 2-Oxocarboxylic acid metabolism | 1 | 19 | 3.44E-01 |
| mmu05221 | Acute myeloid leukemia | 2 | 57 | 3.57E-01 |
| mmu04614 | Renin-angiotensin system | 1 | 20 | 3.58E-01 |
| mmu04713 | Circadian entrainment | 3 | 98 | 3.64E-01 |
| mmu04621 | NOD-like receptor signaling pathway | 2 | 58 | 3.64E-01 |
| mmu04913 | Ovarian steroidogenesis | 2 | 58 | 3.64E-01 |
| mmu04020 | Calcium signaling pathway | 5 | 181 | 3.65E-01 |
| mmu04916 | Melanogenesis | 3 | 100 | 3.76E-01 |
| mmu04310 | Wnt signaling pathway | 4 | 143 | 3.84E-01 |
| mmu04014 | Ras signaling pathway | 6 | 228 | 3.84E-01 |
| mmu04964 | Proximal tubule bicarbonate reclamation | 1 | 22 | 3.86E-01 |
| mmu00562 | Inositol phosphate metabolism | 2 | 61 | 3.88E-01 |
| mmu04150 | mTOR signaling pathway | 2 | 61 | 3.88E-01 |
| mmu04730 | Long-term depression | 2 | 61 | 3.88E-01 |
| mmu04972 | Pancreatic secretion | 3 | 103 | 3.94E-01 |
| mmu00360 | Phenylalanine metabolism | 1 | 23 | 4.00E-01 |
| mmu00601 | Glycosphingolipid biosynthesis - lacto and neolacto series | 1 | 26 | 4.39E-01 |
| mmu00630 | Glyoxylate and dicarboxylate metabolism | 1 | 26 | 4.39E-01 |
| mmu00650 | Butanoate metabolism | 1 | 26 | 4.39E-01 |
| mmu04950 | Maturity onset diabetes of the young | 1 | 26 | 4.39E-01 |
| mmu04622 | RIG-I-like receptor signaling pathway | 2 | 68 | 4.42E-01 |
| mmu03060 | Protein export | 1 | 28 | 4.63E-01 |
| mmu05412 | Arrhythmogenic right ventricular cardiomyopathy (ARVC) | 2 | 74 | 4.85E-01 |
| mmu00514 | Other types of O-glycan biosynthesis | 1 | 31 | 4.98E-01 |
| mmu04670 | Leukocyte transendothelial migration | 3 | 121 | 4.98E-01 |
| mmu03018 | RNA degradation | 2 | 78 | 5.13E-01 |
| mmu04260 | Cardiac muscle contraction | 2 | 78 | 5.13E-01 |
| mmu04810 | Regulation of actin cytoskeleton | 5 | 217 | 5.19E-01 |
| mmu00052 | Galactose metabolism | 1 | 33 | 5.20E-01 |
| mmu04750 | Inflammatory mediator regulation of TRP channels | 3 | 126 | 5.25E-01 |

| | | | | |
|---|---|---|---|---|
| mmu04145 | Phagosome | 4 | 174 | 5.33E-01 |
| mmu04070 | Phosphatidylinositol signaling system | 2 | 81 | 5.34E-01 |
| mmu04210 | Apoptosis | 2 | 82 | 5.40E-01 |
| mmu05323 | Rheumatoid arthritis | 2 | 82 | 5.40E-01 |
| mmu04360 | Axon guidance | 3 | 129 | 5.41E-01 |
| mmu00140 | Steroid hormone biosynthesis | 2 | 86 | 5.66E-01 |
| mmu04911 | Insulin secretion | 2 | 86 | 5.66E-01 |
| mmu05219 | Bladder cancer | 1 | 38 | 5.70E-01 |
| mmu00350 | Tyrosine metabolism | 1 | 39 | 5.80E-01 |
| mmu04727 | GABAergic synapse | 2 | 89 | 5.85E-01 |
| mmu04960 | Aldosterone-regulated sodium reabsorption | 1 | 40 | 5.89E-01 |
| mmu00590 | Arachidonic acid metabolism | 2 | 92 | 6.03E-01 |
| mmu05032 | Morphine addiction | 2 | 92 | 6.03E-01 |
| mmu04910 | Insulin signaling pathway | 3 | 142 | 6.07E-01 |
| mmu04962 | Vasopressin-regulated water reabsorption | 1 | 44 | 6.24E-01 |
| mmu04973 | Carbohydrate digestion and absorption | 1 | 45 | 6.32E-01 |
| mmu04024 | cAMP signaling pathway | 4 | 198 | 6.36E-01 |
| mmu04978 | Mineral absorption | 1 | 46 | 6.40E-01 |
| mmu04064 | NF-kappa B signaling pathway | 2 | 100 | 6.49E-01 |
| mmu04620 | Toll-like receptor signaling pathway | 2 | 101 | 6.54E-01 |
| mmu00600 | Sphingolipid metabolism | 1 | 48 | 6.56E-01 |
| mmu04932 | Non-alcoholic fatty liver disease (NAFLD) | 3 | 154 | 6.61E-01 |
| mmu05168 | Herpes simplex infection | 4 | 205 | 6.63E-01 |
| mmu00510 | N-Glycan biosynthesis | 1 | 49 | 6.64E-01 |
| mmu04330 | Notch signaling pathway | 1 | 49 | 6.64E-01 |
| mmu05030 | Cocaine addiction | 1 | 49 | 6.64E-01 |
| mmu04630 | Jak-STAT signaling pathway | 3 | 155 | 6.66E-01 |
| mmu01212 | Fatty acid metabolism | 1 | 50 | 6.71E-01 |
| mmu03460 | Fanconi anemia pathway | 1 | 52 | 6.85E-01 |
| mmu05014 | Amyotrophic lateral sclerosis (ALS) | 1 | 52 | 6.85E-01 |
| mmu05213 | Endometrial cancer | 1 | 52 | 6.85E-01 |
| mmu00500 | Starch and sucrose metabolism | 1 | 54 | 6.99E-01 |
| mmu05330 | Allograft rejection | 1 | 54 | 6.99E-01 |
| mmu04066 | HIF-1 signaling pathway | 2 | 111 | 7.04E-01 |
| mmu05217 | Basal cell carcinoma | 1 | 55 | 7.06E-01 |

| | | | | |
|---|---|---|---|---|
| mmu03013 | RNA transport | 3 | 166 | 7.10E-01 |
| mmu05206 | MicroRNAs in cancer | 5 | 270 | 7.12E-01 |
| mmu05223 | Non-small cell lung cancer | 1 | 56 | 7.12E-01 |
| mmu04725 | Cholinergic synapse | 2 | 113 | 7.14E-01 |
| mmu04724 | Glutamatergic synapse | 2 | 114 | 7.18E-01 |
| mmu05010 | Alzheimer's disease | 3 | 173 | 7.36E-01 |
| mmu04370 | VEGF signaling pathway | 1 | 60 | 7.37E-01 |
| mmu04940 | Type I diabetes mellitus | 1 | 61 | 7.43E-01 |
| mmu05203 | Viral carcinogenesis | 4 | 229 | 7.45E-01 |
| mmu05152 | Tuberculosis | 3 | 176 | 7.47E-01 |
| mmu04623 | Cytosolic DNA-sensing pathway | 1 | 63 | 7.54E-01 |
| mmu04142 | Lysosome | 2 | 123 | 7.57E-01 |
| mmu04722 | Neurotrophin signaling pathway | 2 | 123 | 7.57E-01 |
| mmu04720 | Long-term potentiation | 1 | 66 | 7.70E-01 |
| mmu05031 | Amphetamine addiction | 1 | 67 | 7.75E-01 |
| mmu04662 | B cell receptor signaling pathway | 1 | 73 | 8.03E-01 |
| mmu04530 | Tight junction | 2 | 136 | 8.04E-01 |
| mmu04917 | Prolactin signaling pathway | 1 | 74 | 8.08E-01 |
| mmu05133 | Pertussis | 1 | 74 | 8.08E-01 |
| mmu05034 | Alcoholism | 3 | 199 | 8.17E-01 |
| mmu05132 | Salmonella infection | 1 | 79 | 8.28E-01 |
| mmu05012 | Parkinson's disease | 2 | 147 | 8.38E-01 |
| mmu03008 | Ribosome biogenesis in eukaryotes | 1 | 82 | 8.39E-01 |
| mmu04912 | GnRH signaling pathway | 1 | 89 | 8.63E-01 |
| mmu00564 | Glycerophospholipid metabolism | 1 | 91 | 8.69E-01 |
| mmu00240 | Pyrimidine metabolism | 1 | 103 | 9.00E-01 |
| mmu04723 | Retrograde endocannabinoid signaling | 1 | 103 | 9.00E-01 |
| mmu04660 | T cell receptor signaling pathway | 1 | 105 | 9.04E-01 |
| mmu04668 | TNF signaling pathway | 1 | 109 | 9.12E-01 |
| mmu05016 | Huntington's disease | 2 | 182 | 9.13E-01 |
| mmu01200 | Carbon metabolism | 1 | 111 | 9.16E-01 |
| mmu04650 | Natural killer cell mediated cytotoxicity | 1 | 119 | 9.30E-01 |
| mmu04062 | Chemokine signaling pathway | 2 | 196 | 9.32E-01 |
| mmu04726 | Serotonergic synapse | 1 | 133 | 9.49E-01 |
| mmu04728 | Dopaminergic synapse | 1 | 133 | 9.49E-01 |

| | | | | |
|---|---|---|---|---|
| mmu05160 | Hepatitis C | 1 | 136 | 9.52E-01 |
| mmu04120 | Ubiquitin mediated proteolysis | 1 | 139 | 9.55E-01 |
| mmu05322 | Systemic lupus erythematosus | 1 | 145 | 9.61E-01 |
| mmu04514 | Cell adhesion molecules (CAMs) | 1 | 160 | 9.72E-01 |
| mmu04080 | Neuroactive ligand-receptor interaction | 1 | 282 | 9.98E-01 |
| mmu04740 | Olfactory transduction | 1 | 1079 | 1.00E+00 |
