## Supplementary material for "An exploration of metabolite and gene responses in mouse skeletal muscles responding to acute sedentary hypoxia": Table S4

| The functional annotation clustering analysis result of cDEGs | | | | | |
|---|---|---|---|---|---|
| ID | Term | p | q | #cDEGs | #total targets |
| Molecular Function | | | | | |
| GO:0005201 | extracellular matrix structural constituent | 6.56E-12 | 6.69E-09 | 15 | 78 |
| GO:0048407 | platelet-derived growth factor binding | 3.89E-10 | 3.97E-07 | 7 | 12 |
| GO:0055131 | C3HC4-type RING finger domain binding | 1.02E-08 | 1.04E-05 | 5 | 6 |
| GO:0019838 | growth factor binding | 2.50E-07 | 2.55E-04 | 14 | 142 |
| GO:0019899 | enzyme binding | 2.88E-06 | 2.93E-03 | 62 | 1929 |
| GO:0019901 | protein kinase binding | 6.67E-06 | 6.80E-03 | 28 | 620 |
| GO:0000982 | transcription factor activity, RNA polymerase II proximal promoter sequence-specific DNA binding | 9.42E-06 | 9.61E-03 | 20 | 365 |
| GO:0005198 | structural molecule activity | 4.28E-05 | 4.37E-02 | 30 | 762 |
| GO:0019900 | kinase binding | 4.72E-05 | 4.81E-02 | 28 | 691 |
| Biological Process | | | | | |
| GO:0030198 | extracellular matrix organization | 1.06E-21 | 5.21E-18 | 41 | 354 |
| GO:0043062 | extracellular structure organization | 1.18E-21 | 5.80E-18 | 41 | 355 |
| GO:0032963 | collagen metabolic process | 4.84E-14 | 2.38E-10 | 20 | 122 |
| GO:0030574 | collagen catabolic process | 6.38E-13 | 3.14E-09 | 15 | 67 |
| GO:0030199 | collagen fibril organization | 7.90E-12 | 3.89E-08 | 12 | 43 |
| GO:0009887 | animal organ morphogenesis | 6.64E-11 | 3.27E-07 | 53 | 1121 |
| GO:0001944 | vasculature development | 1.12E-10 | 5.53E-07 | 39 | 677 |
| GO:0001501 | skeletal system development | 1.16E-10 | 5.70E-07 | 34 | 530 |
| GO:0001568 | blood vessel development | 5.10E-10 | 2.51E-06 | 37 | 651 |
| GO:0072358 | cardiovascular system development | 7.76E-10 | 3.82E-06 | 49 | 1058 |
| GO:0072359 | circulatory system development | 7.76E-10 | 3.82E-06 | 49 | 1058 |
| GO:0050678 | regulation of epithelial cell proliferation | 4.19E-09 | 2.07E-05 | 24 | 323 |
| GO:0048514 | blood vessel morphogenesis | 4.86E-09 | 2.40E-05 | 32 | 551 |
| GO:0010692 | regulation of alkaline phosphatase activity | 6.07E-09 | 2.99E-05 | 6 | 10 |
| GO:1901700 | response to oxygen-containing compound | 1.39E-08 | 6.84E-05 | 61 | 1614 |
| GO:0048646 | anatomical structure formation involved in morphogenesis | 3.14E-08 | 1.55E-04 | 52 | 1299 |
| GO:0050673 | epithelial cell proliferation | 3.99E-08 | 1.97E-04 | 25 | 391 |
| GO:0031399 | regulation of protein modification process | 5.73E-08 | 2.83E-04 | 65 | 1841 |
| GO:0061448 | connective tissue development | 6.98E-08 | 3.44E-04 | 20 | 266 |
| GO:0001503 | ossification | 7.52E-08 | 3.70E-04 | 25 | 404 |
| GO:0001655 | urogenital system development | 1.12E-07 | 5.50E-04 | 23 | 355 |

| | | | | | |
|---|---|---|---|---|---|
| GO:0031400 | negative regulation of protein modification process | 1.25E-07 | 6.14E-04 | 33 | 666 |
| GO:0072001 | renal system development | 1.82E-07 | 8.97E-04 | 21 | 309 |
| GO:0035295 | tube development | 2.46E-07 | 1.21E-03 | 33 | 686 |
| GO:0051241 | negative regulation of multicellular organismal process | 3.24E-07 | 1.59E-03 | 45 | 1127 |
| GO:0010563 | negative regulation of phosphorus metabolic process | 3.84E-07 | 1.89E-03 | 30 | 599 |
| GO:0045936 | negative regulation of phosphate metabolic process | 3.84E-07 | 1.89E-03 | 30 | 599 |
| GO:0048732 | gland development | 4.10E-07 | 2.02E-03 | 27 | 504 |
| GO:0045595 | regulation of cell differentiation | 4.68E-07 | 2.31E-03 | 59 | 1699 |
| GO:0051248 | negative regulation of protein metabolic process | 4.99E-07 | 2.46E-03 | 46 | 1183 |
| GO:0032269 | negative regulation of cellular protein metabolic process | 5.69E-07 | 2.80E-03 | 44 | 1112 |
| GO:0050679 | positive regulation of epithelial cell proliferation | 5.94E-07 | 2.92E-03 | 15 | 175 |
| GO:0048598 | embryonic morphogenesis | 6.53E-07 | 3.22E-03 | 31 | 648 |
| GO:0001933 | negative regulation of protein phosphorylation | 7.58E-07 | 3.74E-03 | 24 | 426 |
| GO:0030857 | negative regulation of epithelial cell differentiation | 7.90E-07 | 3.89E-03 | 8 | 43 |
| GO:0003012 | muscle system process | 8.95E-07 | 4.41E-03 | 24 | 430 |
| GO:0007423 | sensory organ development | 1.02E-06 | 5.00E-03 | 29 | 594 |
| GO:0060429 | epithelium development | 1.11E-06 | 5.45E-03 | 48 | 1296 |
| GO:0051093 | negative regulation of developmental process | 1.12E-06 | 5.50E-03 | 38 | 914 |
| GO:0006805 | xenobiotic metabolic process | 1.12E-06 | 5.52E-03 | 11 | 96 |
| GO:0001822 | kidney development | 1.29E-06 | 6.36E-03 | 19 | 291 |
| GO:2000026 | regulation of multicellular organismal development | 1.65E-06 | 8.15E-03 | 62 | 1893 |
| GO:0071466 | cellular response to xenobiotic stimulus | 1.87E-06 | 9.19E-03 | 11 | 101 |
| GO:0001889 | liver development | 2.12E-06 | 1.05E-02 | 14 | 169 |
| GO:0007167 | enzyme linked receptor protein signaling pathway | 2.21E-06 | 1.09E-02 | 40 | 1016 |
| GO:0009719 | response to endogenous stimulus | 2.22E-06 | 1.09E-02 | 58 | 1740 |
| GO:0009790 | embryo development | 2.41E-06 | 1.19E-02 | 43 | 1135 |
| GO:0009628 | response to abiotic stimulus | 2.55E-06 | 1.26E-02 | 45 | 1216 |
| GO:0061008 | hepaticobiliary system development | 2.61E-06 | 1.29E-02 | 14 | 172 |
| GO:0033993 | response to lipid | 2.62E-06 | 1.29E-02 | 40 | 1023 |
| GO:0045596 | negative regulation of cell differentiation | 2.74E-06 | 1.35E-02 | 31 | 694 |
| GO:0010629 | negative regulation of gene expression | 2.80E-06 | 1.38E-02 | 55 | 1627 |
| GO:0030855 | epithelial cell differentiation | 3.00E-06 | 1.48E-02 | 30 | 662 |
| GO:0045444 | fat cell differentiation | 3.00E-06 | 1.48E-02 | 16 | 225 |
| GO:0001525 | angiogenesis | 3.37E-06 | 1.66E-02 | 24 | 464 |
| GO:0051174 | regulation of phosphorus metabolic process | 3.72E-06 | 1.83E-02 | 58 | 1769 |

| | | | | | |
|---|---|---|---|---|---|
| GO:0009410 | response to xenobiotic stimulus | 3.97E-06 | 1.96E-02 | 11 | 109 |
| GO:0006468 | protein phosphorylation | 4.51E-06 | 2.22E-02 | 62 | 1952 |
| GO:0042326 | negative regulation of phosphorylation | 4.68E-06 | 2.31E-02 | 24 | 473 |
| GO:0016311 | dephosphorylation | 5.48E-06 | 2.70E-02 | 23 | 445 |
| GO:0042127 | regulation of cell proliferation | 5.67E-06 | 2.80E-02 | 55 | 1666 |
| GO:0048568 | embryonic organ development | 5.79E-06 | 2.85E-02 | 24 | 479 |
| GO:0019220 | regulation of phosphate metabolic process | 6.20E-06 | 3.06E-02 | 57 | 1756 |
| GO:0048729 | tissue morphogenesis | 6.77E-06 | 3.34E-02 | 32 | 762 |
| GO:0048562 | embryonic organ morphogenesis | 6.78E-06 | 3.34E-02 | 19 | 326 |
| GO:0023057 | negative regulation of signaling | 7.96E-06 | 3.92E-02 | 46 | 1311 |
| GO:0030856 | regulation of epithelial cell differentiation | 8.01E-06 | 3.95E-02 | 12 | 140 |
| GO:0043408 | regulation of MAPK cascade | 8.71E-06 | 4.29E-02 | 31 | 735 |
| GO:0043409 | negative regulation of MAPK cascade | 9.07E-06 | 4.47E-02 | 13 | 166 |
| GO:0023014 | signal transduction by protein phosphorylation | 9.27E-06 | 4.57E-02 | 37 | 962 |
| GO:0001932 | regulation of protein phosphorylation | 9.85E-06 | 4.85E-02 | 48 | 1404 |
| GO:0007155 | cell adhesion | 1.01E-05 | 4.97E-02 | 51 | 1530 |
| GO:0070848 | response to growth factor | 1.09E-05 | 5.37E-02 | 29 | 671 |
| GO:0010648 | negative regulation of cell communication | 1.11E-05 | 5.46E-02 | 46 | 1328 |
| GO:0043067 | regulation of programmed cell death | 1.12E-05 | 5.53E-02 | 51 | 1536 |
| GO:0070482 | response to oxygen levels | 1.13E-05 | 5.58E-02 | 19 | 338 |
| GO:0070208 | protein heterotrimerization | 1.24E-05 | 6.10E-02 | 5 | 18 |
| GO:0051270 | regulation of cellular component movement | 1.24E-05 | 6.10E-02 | 34 | 860 |
| GO:0022610 | biological adhesion | 1.25E-05 | 6.15E-02 | 51 | 1542 |
| GO:0001101 | response to acid chemical | 1.28E-05 | 6.31E-02 | 20 | 372 |
| GO:0071363 | cellular response to growth factor stimulus | 1.36E-05 | 6.67E-02 | 28 | 643 |
| GO:1901698 | response to nitrogen compound | 1.43E-05 | 7.03E-02 | 37 | 981 |
| GO:0006470 | protein dephosphorylation | 1.45E-05 | 7.15E-02 | 19 | 344 |
| GO:0006936 | muscle contraction | 1.70E-05 | 8.39E-02 | 19 | 348 |
| GO:0048585 | negative regulation of response to stimulus | 1.72E-05 | 8.45E-02 | 50 | 1518 |
| GO:0042981 | regulation of apoptotic process | 1.75E-05 | 8.60E-02 | 50 | 1519 |
| **Cellular Component** | | | | | |
| GO:0031012 | extracellular matrix | 1.85E-18 | 8.70E-16 | 41 | 444 |
| GO:0044420 | extracellular matrix component | 2.32E-18 | 1.10E-15 | 25 | 141 |
| GO:0098644 | complex of collagen trimers | 4.12E-14 | 1.94E-11 | 11 | 23 |
| GO:0005788 | endoplasmic reticulum lumen | 1.70E-12 | 8.02E-10 | 23 | 207 |

| | | | | | |
|---|---|---|---|---|---|
| GO:0005581 | collagen trimer | 2.90E-11 | 1.37E-08 | 15 | 88 |
| GO:0098643 | banded collagen fibril | 3.29E-10 | 1.55E-07 | 7 | 12 |
| GO:0005583 | fibrillar collagen trimer | 3.29E-10 | 1.55E-07 | 7 | 12 |
| GO:0005604 | basement membrane | 6.68E-10 | 3.15E-07 | 15 | 109 |
| GO:0005615 | extracellular space | 1.66E-09 | 7.81E-07 | 58 | 1449 |
| GO:0005588 | collagen type V trimer | 5.28E-06 | 2.49E-03 | 3 | 3 |
| GO:0031983 | vesicle lumen | 1.12E-04 | 5.26E-02 | 9 | 108 |
| **Domain** | | | | | |
| PF01391 | Collagen | 2.19E-11 | 3.96E-08 | 15 | 85 |
| IPR008160 | Collagen | 2.19E-11 | 3.96E-08 | 15 | 85 |
| PS51461 | NC1_FIB | 1.57E-10 | 2.83E-07 | 7 | 11 |
| IPR000885 | Fib_collagen_C | 1.57E-10 | 2.83E-07 | 7 | 11 |
| SM00038 | COLFI | 1.57E-10 | 2.83E-07 | 7 | 11 |
| PD002078 | Fib_collagen_C | 1.57E-10 | 2.83E-07 | 7 | 11 |
| PF01410 | COLFI | 1.57E-10 | 2.83E-07 | 7 | 11 |
| SM00282 | LamG | 8.89E-07 | 1.61E-03 | 8 | 44 |
| PF02210 | Laminin_G_2 | 5.70E-06 | 1.03E-02 | 7 | 40 |
| PF00012 | HSP70 | 6.22E-06 | 1.12E-02 | 5 | 16 |
| IPR001791 | Laminin_G | 7.75E-06 | 1.40E-02 | 8 | 58 |
| PS01036 | HSP70_3 | 8.68E-06 | 1.57E-02 | 5 | 17 |
| PS00297 | HSP70_1 | 8.68E-06 | 1.57E-02 | 5 | 17 |
| PS00329 | HSP70_2 | 8.68E-06 | 1.57E-02 | 5 | 17 |
| IPR013126 | Hsp_70_fam | 1.18E-05 | 2.14E-02 | 5 | 18 |
| PS01286 | FA58C_2 | 2.69E-05 | 4.87E-02 | 5 | 21 |
| PS01285 | FA58C_1 | 2.69E-05 | 4.87E-02 | 5 | 21 |
| PS50022 | FA58C_3 | 2.69E-05 | 4.87E-02 | 5 | 21 |
| SM00231 | FA58C | 2.69E-05 | 4.87E-02 | 5 | 21 |
| IPR029047 | HSP70_peptide-bd | 4.23E-05 | 7.65E-02 | 4 | 12 |
| IPR000421 | FA58C | 5.39E-05 | 9.73E-02 | 5 | 24 |
| PF00754 | F5_F8_type_C | 5.39E-05 | 9.73E-02 | 5 | 24 |
| **Gene Family** | | | | | |
| 490 | Collagens | 1.25E-14 | 1.97E-12 | 13 | 46 |
| 583 | Heat shock 70kDa proteins | 1.80E-06 | 2.85E-04 | 5 | 17 |
| 626 | Laminin subunits | 1.20E-05 | 1.89E-03 | 4 | 12 |
