## Supplementary material for "An exploration of metabolite and gene responses in mouse skeletal muscles responding to acute sedentary hypoxia": Table S5

| Metabolic reactions containing cDEGs in the genome-based mouse metabolic network | | | | | | |
|---|---|---|---|---|---|---|
| **Reaction ID** | **Reaction Name** | **Reaction** | **Subsystem** | **Compartment** | **Genes** | **Folds** |
| 2HCO3_NAt | Bicarbonate transport (Na/HCO3 1:2 cotransport) | na1_e + 2.0 hco3_e <-> 2.0 hco3_c + na1_c | Transport, Extracellular | cytosol | Slc4a4 | 0.55556 |
| 34DHALDD | Aldehyde dehydrogenase 3 4 dihydroxyphenylacetaldehyde NAD | 34dhpac_c + h2o_c + nad_c <-> nadh_c + 2.0 h_c + 34dhpha_c | Tyrosine metabolism | cytosol | Aldh3a1 | 14.95 |
| 34DHPLACOX_NADP | 34DHPLACOX LPAREN NADP RPAREN | h2o_c + 34dhpac_c + nadp_c <-> nadph_c + 2.0 h_c + 34dhpha_c | Tyrosine metabolism | cytosol | Aldh3a1 | 14.95 |
| 34DHXMANDACOX | 3,4-Dihydroxymandelaldehyde:NAD+ oxidoreductase | h2o_c + 34dhmald_c + nad_c <-> 34dhoxmand_c + nadh_c + 2.0 h_c | Tyrosine metabolism | cytosol | Aldh3a1 | 14.95 |
| 3HCO3_NAt | Bicarbonate transport (Na/HCO3 1:3 cotransport) | na1_e + 3.0 hco3_e <-> 3.0 hco3_c + na1_c | Transport, Extracellular | cytosol | Slc4a4 | 0.55556 |
| 3M4HDXPAC | 3-Methoxy-4-hydroxyphenylacetaldehyde:NAD+ oxidoreductase | 3mox4hpac_c + nad_c + h2o_c <-> homoval_c + nadh_c + 2.0 h_c | Tyrosine metabolism | cytosol | Aldh3a1 | 14.95 |
| 3MOX4HOXPGALDOX | 3-Methoxy-4-hydroxyphenylglycolaldehyde: NAD+ oxidoreductase | h2o_c + nad_c + 3m4hpga_c <-> 3mox4hoxm_c + nadh_c + 2.0 h_c | Tyrosine metabolism | cytosol | Aldh3a1 | 14.95 |
| 3MOX4HOXPGALDOX_NADP | 3-Methoxy-4-hydroxyphenylglycolaldehyde: NADP+ oxidoreductase | h2o_c + 3m4hpga_c + nadp_c <-> nadph_c + 3mox4hoxm_c + 2.0 h_c | Tyrosine metabolism | cytosol | Aldh3a1 | 14.95 |
| 4HOXPACDOX_NADP | 4HOXPACDOX LPAREN NADP RPAREN | h2o_c + 4hoxpacd_c + nadp_c <-> 4hphac_c + nadph_c + 2.0 h_c | Tyrosine metabolism | cytosol | Aldh3a1 | 14.95 |
| 5HOXINDACTOX | 5-Hydroxyindoleacetaldehyde:NAD+ oxidoreductase | h2o_c + 5hoxindact_c + nad_c <-> 5hoxindoa_c + nadh_c + 2.0 h_c | Tryptophan metabolism | cytosol | Aldh9a1 | 0.52356 |
| ABUTD | Aminobutyraldehyde dehydrogenase | 4abutn_c + h2o_c + nad_c <-> 4abut_c + 2.0 h_c + nadh_c | beta-Alanine metabolism | cytosol | Aldh9a1 | 0.52356 |
| ADNCYC | Adenylate cyclase | atp_c <-> camp_c + ppi_c | Nucleotides | cytosol | Adcy7 | 0.52083 |

| | | | | | | |
|---|---|---|---|---|---|---|
| ALAt4 | Alanine-Sodium symporter | ala__L_e + na1_e <-> ala__L_c + na1_c | Transport, Extracellular | cytosol | Slc38a4 | 0.52356 |
| ALDD19x_P | Aldehyde dehydrogenase (phenylacetaldehyde, NADP) | pacald_c + nadp_c + h2o_c <-> nadph_c + pac_c + 2.0 h_c | Phenylalanine metabolism | cytosol | Aldh3a1 | 14.95 |
| ALDD19xr | Aldehyde dehydrogenase (phenylacetaldehyde, NAD) | h2o_c + nad_c + pacald_c <-> 2.0 h_c + nadh_c + pac_c | Phenylalanine metabolism | cytosol | Aldh3a1 | 14.95 |
| ALDD20x | Aldehyde dehydrogenase (indole-3-acetaldehyde, NAD) | nad_c + h2o_c + id3acald_c <-> 2.0 h_c + nadh_c + ind3ac_c | Tryptophan metabolism | cytosol | Aldh9a1 | 0.52356 |
| ALDD2x | Aldehyde dehydrogenase (acetaldehyde, NAD) | acald_c + h2o_c + nad_c <-> ac_c + 2.0 h_c + nadh_c | Glycolysis/Gluconeogenesis | cytosol | Aldh3a1; Aldh9a1 | 14.95;0.52356 |
| ALDD2y | Aldehyde dehydrogenase (acetaldehyde, NADP) | acald_c + h2o_c + nadp_c <-> ac_c + 2.0 h_c + nadph_c | Glycolysis/Gluconeogenesis | cytosol | Aldh3a1; Aldh9a1 | 14.95;0.52356 |
| ARGt5r | ARGtiDF | arg__L_e <-> arg__L_c | Transport, Extracellular | cytosol | Slc38a4 | 0.52356 |
| ASNt4 | L-asparagine transport in via sodium symport | na1_e + asn__L_e <-> asn__L_c + na1_c | Transport, Extracellular | cytosol | Slc38a4 | 0.52356 |
| BAMPPALDOX | Beta-Aminopropion aldehyde NAD+ oxidoreductase | nad_c + h2o_c + bamppald_c <-> 2.0 h_c + nadh_c + ala_B_c | beta-Alanine metabolism | cytosol | Aldh9a1 | 0.52356 |
| BETALDHx | Betaine-aldehyde dehydrogenase | betald_c + h2o_c + nad_c <-> glyb_c + 2.0 h_c + nadh_c | Glycine, Serine, and Threonine Metabolism | cytosol | Aldh9a1 | 0.52356 |
| CGLYt3_2 | Cys-Gly transport in via proton symport | 2.0 h_e + cgly_e <-> cgly_c + 2.0 h_c | Transport, Extracellular | cytosol | Slc15a2 | 5.26 |
| DHPM1 | Dihydropyrimidinase (5,6-dihydrouracil) | h2o_c + 56dura_c <-> cala_c + h_c | Pyrimidine Catabolism | cytosol | Dpysl3 | 0.40984 |
| DOPAtu | Dopamine uniport | dopa_e <-> dopa_c | Transport, Extracellular | cytosol | Slc22a3 | 0.17331 |
| ESTSULT | Estrogen sulfotransferase | paps_c + estrone_c <-> pap_c + h_c + estrones_c | Steroid Metabolism | cytosol | Sult1a1 | 1.62 |
| FACOAL150 | Fatty-acid--CoA ligase | atp_c + coa_c + ptdca_c <-> ppi_c + amp_c + ptdcacoa_c | Fatty acid activation | cytosol | Acsl1 | 0.55249 |
| FACOAL160 | Fatty acid CoA ligase hexadecanoate | atp_c + coa_c + hdca_c <-> amp_c + pmtcoa_c + ppi_c | Fatty acid activation | cytosol | Acsl1 | 0.55249 |

| | | | | | | |
|---|---|---|---|---|---|---|
| FACOAL161 | Fatty acid CoA ligase hexadecenoate | atp_c + coa_c + hdcea_c <-> amp_c + hdcoa_c + ppi_c | Fatty acid activation | cytosol | Acsl1 | 0.55249 |
| FACOAL170 | Fatty-acid--CoA ligase | coa_c + hpdca_c + atp_c <-> hpdcacoa_c + ppi_c + amp_c | Fatty acid activation | cytosol | Acsl1 | 0.55249 |
| FACOAL180 | Fatty acid CoA ligase octadecanoate | atp_c + coa_c + ocdca_c <-> amp_c + ppi_c + stcoa_c | Fatty acid activation | cytosol | Acsl1 | 0.55249 |
| FACOAL181 | Fatty acid CoA ligase octadecenoate | atp_c + coa_c + ocdcea_c <-> amp_c + odecoa_c + ppi_c | Fatty acid activation | cytosol | Acsl1 | 0.55249 |
| FACOAL1812 | Fatty-acid--CoA ligase | coa_c + vacc_c + atp_c <-> vacccoa_c + ppi_c + amp_c | Fatty acid activation | cytosol | Acsl1 | 0.55249 |
| FACOAL1813 | Fatty-acid--CoA ligase | atp_c + coa_c + elaid_c <-> od2coa_c + ppi_c + amp_c | Fatty acid activation | cytosol | Acsl1 | 0.55249 |
| FACOAL1821 | Linoleic acid:CoA ligase (AMP-forming) | atp_c + coa_c + lnlc_c <-> lnlccoa_c + ppi_c + amp_c | Fatty acid activation | cytosol | Acsl1 | 0.55249 |
| FACOAL1822 | Fatty-acid--CoA ligase | lneldc_c + atp_c + coa_c <-> lneldccoa_c + ppi_c + amp_c | Fatty acid activation | cytosol | Acsl1 | 0.55249 |
| FACOAL1831 | Gamma-Linolenic acid:CoA ligase (AMP-forming) | atp_c + coa_c + lnlncg_c <-> ppi_c + lnlncgcoa_c + amp_c | Fatty acid activation | cytosol | Acsl1 | 0.55249 |
| FACOAL1832 | Alpha-Linolenic acid:CoA ligase (AMP-forming) | coa_c + lnlnca_c + atp_c <-> lnlncacoa_c + ppi_c + amp_c | Fatty acid activation | cytosol | Acsl1 | 0.55249 |
| FACOAL184 | Fatty-acid--CoA ligase | strdnc_c + atp_c + coa_c <-> strdnccoa_c + ppi_c + amp_c | Fatty acid activation | cytosol | Acsl1 | 0.55249 |
| FACOAL200 | Fatty-acid--CoA ligase | atp_c + arach_c + coa_c <-> amp_c + arachcoa_c + ppi_c | Fatty acid activation | cytosol | Acsl1 | 0.55249 |
| FACOAL203 | Fatty-acid--CoA ligase | atp_c + coa_c + dlnlcg_c <-> ppi_c + dlnlcgcoa_c + amp_c | Fatty acid activation | cytosol | Acsl1 | 0.55249 |
| FACOAL204 | Fatty-acid--CoA ligase | coa_c + arachd_c + atp_c <-> arachdcoa_c + ppi_c + amp_c | Fatty acid activation | cytosol | Acsl1 | 0.55249 |
| FACOAL204 | Fatty-acid--CoA ligase | coa_c + arachd_c + atp_c <-> arachdcoa_c + ppi_c + amp_c | Fatty acid activation | cytosol | Acsl1 | 0.55249 |
| FACOAL2042 | Fatty-acid--CoA ligase | eicostet_c + atp_c + coa_c <-> eicostetcoa_c + ppi_c + amp_c | Fatty acid activation | cytosol | Acsl1 | 0.55249 |
| FACOAL205 | Fatty-acid--CoA ligase | coa_c + tmndnc_c + atp_c <-> tmndnccoa_c + ppi_c + amp_c | Fatty acid activation | cytosol | Acsl1 | 0.55249 |

| | | | | | | |
|---|---|---|---|---|---|---|
| FACOAL224 | Fatty-acid--CoA ligase | coa_c + adrn_c + atp_c <-> adrncoa_c + ppi_c + amp_c | Fatty acid activation | cytosol | Acsl1 | 0.55249 |
| FACOAL2251 | Fatty-acid--CoA ligase | atp_c + coa_c + dcsptn1_c <-> dcsptn1coa_c + ppi_c + amp_c | Fatty acid activation | cytosol | Acsl1 | 0.55249 |
| FACOAL2252 | Fatty-acid--CoA ligase | clpnd_c + coa_c + atp_c <-> clpndcoa_c + ppi_c + amp_c | Fatty acid activation | cytosol | Acsl1 | 0.55249 |
| FACOAL226 | Fatty-acid--CoA ligase | coa_c + atp_c + crvnc_c <-> c226coa_c + amp_c + ppi_c | Fatty acid activation | cytosol | Acsl1 | 0.55249 |
| FACOAL240 | Fatty-acid--CoA ligase | atp_c + coa_c + lgnc_c <-> ppi_c + amp_c + lgnccoa_c | Fatty acid activation | cytosol | Acsl1 | 0.55249 |
| FACOAL241 | Fatty-acid--CoA ligase | nrvnc_c + atp_c + coa_c <-> nrvnccoa_c + ppi_c + amp_c | Fatty acid activation | cytosol | Acsl1 | 0.55249 |
| FACOAL244_1 | Fatty-acid--CoA ligase | atp_c + coa_c + tettet6_c <-> tettet6coa_c + ppi_c + amp_c | Fatty acid activation | cytosol | Acsl1 | 0.55249 |
| FACOAL245_1 | Fatty-acid--CoA ligase | atp_c + coa_c + tetpent6_c <-> ppi_c + amp_c + tetpent6coa_c | Fatty acid activation | cytosol | Acsl1 | 0.55249 |
| FACOAL245_2 | Fatty-acid--CoA ligase | tetpent3_c + atp_c + coa_c <-> tetpent3coa_c + ppi_c + amp_c | Fatty acid activation | cytosol | Acsl1 | 0.55249 |
| FACOAL246_1 | Fatty-acid--CoA ligase | atp_c + coa_c + tethex3_c <-> tethex3coa_c + ppi_c + amp_c | Fatty acid activation | cytosol | Acsl1 | 0.55249 |
| FACOAL260 | Fatty-acid--CoA ligase (n-C26:0) | atp_c + coa_c + hexc_c <-> hexccoa_c + ppi_c + amp_c | Fatty acid activation | cytosol | Acsl1 | 0.55249 |
| G3PD2m | Glycerol-3-phosphate dehydrogenase (FAD), mitochondrial | fad_m + glyc3p_c <-> dhap_c + fadh2_m | Glycolysis/Gluconeogenesis | cytosol | Gpd2 | 1.44 |
| GCALDD | Glycolaldehyde dehydrogenase | gcald_c + h2o_c + nad_c <-> glyclt_c + 2.0 h_c + nadh_c | Glyoxylate and Dicarboxylate Metabolism | cytosol | Aldh3a1; Aldh9a1 | 14.95;0.52356 |
| GGNG | Glycogenin self-glucosylation | Tyr_ggn_c + 8.0 udpg_c <-> 8.0 udp_c + ggn_c + 8.0 h_c | Starch and Sucrose Metabolism | cytosol | Gyg | 1.55 |
| GLACO | D-Glucuronolactone:NAD+ oxidoreductase | 2.0 h2o_c + glac_c + nad_c <-> nadh_c + 3.0 h_c + glcr_c | Ascorbate and Aldarate Metabolism | cytosol | Aldh3a1; Aldh9a1 | 14.95;0.52356 |

| | | | | | | |
|---|---|---|---|---|---|---|
| GLGNS1 | Glycogen synthase (ggn -> glygn1) | ggn_c + 3.0 udpg_c <-> 3.0 udp_c + 3.0 h_c + glygn1_c | Starch and Sucrose Metabolism | cytosol | Gyg | 1.55 |
| GLNS | Glutamine synthetase | atp_c + glu__L_c + nh4_c <-> adp_c + gln__L_c + h_c + pi_c | Glutamate metabolism | cytosol | Glul | 1.82 |
| GLNt4 | L-glutamine reversible transport via sodium symport | na1_e + gln__L_e <-> gln__L_c + na1_c | Transport, Extracellular | cytosol | Slc38a4 | 0.52356 |
| GLYAMDTRc | Glycine amidinotransferase (c) | gly_c + arg__L_c <-> orn_c + gudac_c | Urea cycle/amino group metabolism | cytosol | Gatm | 0.60976 |
| GLYt4 | Glycine transport via sodium symport | na1_e + gly_e <-> gly_c + na1_c | Transport, Extracellular | cytosol | Slc38a4 | 0.52356 |
| GNMT | Glycine N-methyltransferase | amet_c + gly_c <-> sarcs_c + h_c + ahcys_c | Glycine, Serine, and Threonine Metabolism | cytosol | Gnmt | 2.87 |
| GTHPe | Glutathione peroxidase | 2.0 gthrd_e + h2o2_e <-> gthox_e + 2.0 h2o_e | Glutathione Metabolism | cytosol | Gpx3 | 1.78 |
| HISTAtu | Histamine uniport | hista_e <-> hista_c | Transport, Extracellular | cytosol | Slc22a3 | 0.17331 |
| HIStiDF | L-histidine transport via diffusion (extracellular to cytosol) | his__L_e <-> his__L_c | Transport, Extracellular | cytosol | Slc38a4 | 0.52356 |
| HSD17B1 | Testicular 17-beta-hydroxysteroid dehydrogenase | nadph_c + h_c + estrone_c <-> nadp_c + estradiol_c | Steroid Metabolism | cytosol | H2-Ke6 | 1.55 |
| IMACTD | Imidazole acetaldeyde dehydrogenase | h2o_c + im4act_c + nad_c <-> 2.0 h_c + im4ac_c + nadh_c | Histidine Metabolism | cytosol | Aldh9a1 | 0.52356 |
| KCC2t | K+-Cl- cotransporter (NH4+) | cl_e + nh4_e <-> cl_c + nh4_c | Transport, Extracellular | cytosol | Slc12a4 | 1.49 |
| KCCt | K+-Cl- cotransport | cl_e + k_e <-> cl_c + k_c | Transport, Extracellular | cytosol | Slc12a4 | 1.49 |
| LCADi | Lactaldehyde dehydrogenase | h2o_c + lald__L_c + nad_c <-> 2.0 h_c + lac__L_c + nadh_c | Pyruvate Metabolism | cytosol | Aldh3a1; Aldh9a1 | 14.95;0.52356 |
| LCADi_D | Lactaldehyde dehydrogenase | lald__D_c + nad_c + h2o_c <-> nadh_c + lac__D_c + 2.0 h_c | Pyruvate Metabolism | cytosol | Aldh3a1; Aldh9a1 | 14.95;0.52356 |
| LEUt4 | L-leucine transport in via sodium symport | na1_e + leu__L_e <-> leu__L_c + na1_c | Transport, Extracellular | cytosol | Slc38a4 | 0.52356 |

| | | | | | | |
|---|---|---|---|---|---|---|
| LYSt5r | LYStiDF | lys__L_e <-> lys__L_c | Transport, Extracellular | cytosol | Slc38a4 | 0.52356 |
| MACOXO | 3-Methylimidazole acetaldehyde:NAD+ oxidoreductase | h2o_c + nad_c + 3mldz_c <-> 2.0 h_c + nadh_c + 3mlda_c | Histidine Metabolism | cytosol | Aldh3a1 | 14.95 |
| MCPST | 3-mercaptopyruvate sulfurtransferase | cyan_c + mercppyr_c <-> h_c + pyr_c + tcynt_c | Cysteine Metabolism | cytosol | Mpst | 1.57 |
| MI1PP | Myo-inositol 1-phosphatase | h2o_c + mi1p__D_c <-> inost_c + pi_c | Inositol Phosphate Metabolism | cytosol | Impa2 | 0.43103 |
| MI3PP | 1D-myo-Inositol 3-phosphate phosphohydrolase | mi3p__D_c + h2o_c <-> inost_c + pi_c | Inositol Phosphate Metabolism | cytosol | Impa2 | 0.43103 |
| MI4PP | 1D-myo-Inositol 4-phosphate phosphohydrolase | mi4p__D_c + h2o_c <-> inost_c + pi_c | Inositol Phosphate Metabolism | cytosol | Impa2 | 0.43103 |
| NABTNO | N4 AcetylaminobutanalNAD oxidoreductase | h2o_c + n4abutn_c + nad_c <-> 4aabutn_c + 2.0 h_c + nadh_c | Arginine and Proline Metabolism | cytosol | Aldh9a1 | 0.52356 |
| NRPPHRtu | Norepinephrine uniport | nrpphr_e <-> nrpphr_c | Transport, Extracellular | cytosol | Slc22a3 | 0.17331 |
| NTD2 | 5'-nucleotidase (UMP) | h2o_c + ump_c <-> pi_c + uri_c | Pyrimidine Catabolism | cytosol | Nt5c3 | 0.59524 |
| NTD4 | 5'-nucleotidase (CMP) | cmp_c + h2o_c <-> cytd_c + pi_c | Pyrimidine Catabolism | cytosol | Nt5c3 | 0.59524 |
| ORNDC | Ornithine Decarboxylase | h_c + orn_c <-> co2_c + ptrc_c | Urea cycle/amino group metabolism | cytosol | Odc1 | 0.5 |
| P4502D6 | Cytochrome P450 2D6 | nadph_c + debrisoquine_c + o2_c + h_c <-> h2o_c + 4hdebrisoquine_c + nadp_c | CYP Metabolism | cytosol | Cyp2d22 | 1.72 |
| PDE1 | 3',5'-cyclic-nucleotide phosphodiesterase | camp_c + h2o_c <-> amp_c + h_c | Nucleotides | cytosol | Pde10a | 0.36765 |
| PDE4 | 3',5'-cyclic-nucleotide phosphodiesterase | 35cgmp_c + h2o_c <-> gmp_c + h_c | Nucleotides | cytosol | Pde10a | 0.36765 |
| PHEt4 | L-phenylalanine transport in via sodium symport | phe__L_e + na1_e <-> phe__L_c + na1_c | Transport, Extracellular | cytosol | Slc38a4 | 0.52356 |
| PROt4 | Na+/Proline-L symporter | na1_e + pro__L_e <-> na1_c + pro__L_c | Transport, Extracellular | cytosol | Slc38a4 | 0.52356 |

| | | | | | | |
|---|---|---|---|---|---|---|
| PTHPS | 6-pyruvoyltetrahydropterin synthase | ahdt_c <-> 6pthp_c + pppi_c | Tetrahydrobiopterin | cytosol | Pts | 1.64 |
| PYLALDOX | Perillyl aldehyde:NAD+ oxidoreductase | pylald_c + nad_c + h2o_c <-> peracd_c + nadh_c + 2.0 h_c | Limonene and pinene degradation | cytosol | Aldh9a1 | 0.52356 |
| SERt4 | L-serine via sodium symport | na1_e + ser__L_e <-> na1_c + ser__L_c | Transport, Extracellular | cytosol | Slc38a4 | 0.52356 |
| SIAASE | SIAASE | s2l2n2m2mn_c + 2.0 h2o_c <-> l2n2m2mn_c + 2.0 acnam_c | N-Glycan Degradation | cytosol | Neu2 | 1.56 |
| SRTNMTX | S-Adenosyl-L-methionine:amine N-methyltransferase (srtn) | amet_c + srtn_c <-> nmthsrtn_c + ahcys_c + h_c | Tryptophan metabolism | cytosol | Inmt | 3.98 |
| TMABADH | 4-trimethylaminobutyraldehyde dehydrogenase | nad_c + 4tmeabut_c + h2o_c <-> 2.0 h_c + 4tmeabutn_c + nadh_c | Lysine Metabolism | cytosol | Aldh9a1 | 0.52356 |
| TRIODTHYSULT | Triiodothyronine Sulfotransferase | triodthy_c + paps_c <-> pap_c + triodthysuf_c + h_c | Tyrosine metabolism | cytosol | Sult1a1 | 1.62 |
| XYLUR | Xylulose reductase | nadph_c + h_c + xylu__L_c <-> xylt_c + nadp_c | Pentose and Glucuronate Interconversions | cytosol | Dcxr | 1.73 |
| ADPRIBt | ADPribose transport | adprib_e <-> adprib_c | Transport, Extracellular | extracellular | Cd38 | 0.54054 |
| AMY1e | Alpha-amylase, extracellular (strch1 -> strch2) | 8.0 h2o_e + strch1_e <-> strch2_e + 8.0 glc__D_e | Starch and Sucrose Metabolism | extracellular | Amy1 | 1.98 |
| AMY2e | Alpha-amylase, extracellular (glygn2 -> glygn4) | 8.0 h2o_e + glygn2_e <-> glygn4_e + 8.0 glc__D_e | Starch and Sucrose Metabolism | extracellular | Amy1 | 1.98 |
| NADNe | NAD nucleosidase,extracellular | h2o_e + nad_e <-> ncam_e + adprib_e + h_e | NAD Metabolism | extracellular | Cd38 | 0.54054 |
| RAI3 | 13-cis-retinoic acid isomerase | retn_c <-> 13_cis_retn_c | Vitamin A Metabolism | extracellular | Gstm2 | 3.64 |
| G14T10g | Beta-N-acetylglucosaminylglycopeptide beta-1,4-galactosyltransferase, Golgi | udpgal_g + ksi_pre14_g <-> h_g + udp_g + ksi_pre15_g | Keratan sulfate biosynthesis | golgi | B4galt1; B4galt5 | 0.55866;0.41667 |

| | | | | | | |
|---|---|---|---|---|---|---|
| G14T11g | Beta-N-acetylglucosaminylglycopeptide beta-1,4-galactosyltransferase, Golgi | udpgal_g + ksi_pre17_g <-> h_g + udp_g + ksi_pre18_g | Keratan sulfate biosynthesis | golgi | B4galt1; B4galt5 | 0.55866;0.41667 |
| G14T12g | Beta-N-acetylglucosaminylglycopeptide beta-1,4-galactosyltransferase, Golgi | udpgal_g + ksi_pre20_g <-> h_g + udp_g + ksi_pre21_g | Keratan sulfate biosynthesis | golgi | B4galt1; B4galt5 | 0.55866;0.41667 |
| G14T13g | Beta-N-acetylglucosaminylglycopeptide beta-1,4-galactosyltransferase, Golgi | udpgal_g + ksi_pre23_g <-> h_g + udp_g + ksi_pre24_g | Keratan sulfate biosynthesis | golgi | B4galt1; B4galt5 | 0.55866;0.41667 |
| G14T14g | Beta-N-acetylglucosaminylglycopeptide beta-1,4-galactosyltransferase, Golgi | udpgal_g + ksi_pre26_g <-> h_g + udp_g + ksi_pre27_g | Keratan sulfate biosynthesis | golgi | B4galt1; B4galt5 | 0.55866;0.41667 |
| G14T15g | Beta-N-acetylglucosaminylglycopeptide beta-1,4-galactosyltransferase, Golgi | udpgal_g + ksi_pre29_g <-> h_g + udp_g + ksi_pre30_g | Keratan sulfate biosynthesis | golgi | B4galt1; B4galt5 | 0.55866;0.41667 |
| G14T16g | Beta-N-acetylglucosaminylglycopeptide beta-1,4-galactosyltransferase, Golgi | udpgal_g + ksi_pre32_g <-> h_g + udp_g + ksi_pre33_g | Keratan sulfate biosynthesis | golgi | B4galt1; B4galt5 | 0.55866;0.41667 |
| G14T17g | Beta-N-acetylglucosaminylglycopeptide beta-1,4-galactosyltransferase, Golgi | udpgal_g + ksi_pre35_g <-> h_g + udp_g + ksi_pre36_g | Keratan sulfate biosynthesis | golgi | B4galt1; B4galt5 | 0.55866;0.41667 |
| G14T18g | Beta-N-acetylglucosaminylglycopeptide beta-1,4-galactosyltransferase, Golgi | 2.0 udpgal_g + core4_g <-> 2.0 h_g + 2.0 udp_g + ksii_core4_pre1_g | Keratan sulfate biosynthesis | golgi | B4galt1; B4galt5 | 0.55866;0.41667 |

| | | | | | | |
|---|---|---|---|---|---|---|
| G14T19g | Beta-N-acetylglucosaminylglycopeptide beta-1,4-galactosyltransferase, Golgi | udpgal_g + ksii_core4_pre3_g <-> h_g + udp_g + ksii_core4_pre4_g | Keratan sulfate biosynthesis | golgi | B4galt1; B4galt5 | 0.55866;0.41667 |
| G14T20g | Beta-N-acetylglucosaminylglycopeptide beta-1,4-galactosyltransferase, Golgi | udpgal_g + ksii_core4_pre6_g <-> h_g + udp_g + ksii_core4_pre7_g | Keratan sulfate biosynthesis | golgi | B4galt1; B4galt5 | 0.55866;0.41667 |
| G14T21g | Beta-N-acetylglucosaminylglycopeptide beta-1,4-galactosyltransferase, Golgi | udpgal_g + ksii_core4_pre9_g <-> h_g + udp_g + ksii_core4_pre10_g | Keratan sulfate biosynthesis | golgi | B4galt1; B4galt5 | 0.55866;0.41667 |
| G14T2g | Beta-N-acetylglucosaminylglycopeptide beta-1,4-galactosyltransferase, Golgi | udpgal_g + ksii_core2_pre1_g <-> h_g + ksii_core2_pre2_g + udp_g | Keratan sulfate biosynthesis | golgi | B4galt1; B4galt5 | 0.55866;0.41667 |
| G14T3g | Beta-N-acetylglucosaminylglycopeptide beta-1,4-galactosyltransferase, Golgi | udpgal_g + ksii_core2_pre3_g <-> h_g + udp_g + ksii_core2_pre4_g | Keratan sulfate biosynthesis | golgi | B4galt1; B4galt5 | 0.55866;0.41667 |
| G14T4g | Beta-N-acetylglucosaminylglycopeptide beta-1,4-galactosyltransferase, Golgi | udpgal_g + ksii_core2_pre6_g <-> h_g + udp_g + ksii_core2_pre7_g | Keratan sulfate biosynthesis | golgi | B4galt1; B4galt5 | 0.55866;0.41667 |
| G14T5g | Beta-N-acetylglucosaminylglycopeptide beta-1,4-galactosyltransferase, Golgi | udpgal_g + ksii_core2_pre9_g <-> h_g + udp_g + ksii_core2_pre10_g | Keratan sulfate biosynthesis | golgi | B4galt1; B4galt5 | 0.55866;0.41667 |
| G14T6g | Beta-N-acetylglucosaminylglycopeptide beta-1,4-galactosyltransferase, Golgi | udpgal_g + ksi_pre2_g <-> h_g + udp_g + ksi_pre3_g | Keratan sulfate biosynthesis | golgi | B4galt1; B4galt5 | 0.55866;0.41667 |

| | | | | | | |
|---|---|---|---|---|---|---|
| G14T7g | Beta-N-acetylglucosaminylglycopeptide beta-1,4-galactosyltransferase, Golgi | udpgal_g + ksi_pre5_g <-> h_g + udp_g + ksi_pre6_g | Keratan sulfate biosynthesis | golgi | B4galt1; B4galt5 | 0.55866;0.41667 |
| G14T8g | Beta-N-acetylglucosaminylglycopeptide beta-1,4-galactosyltransferase, Golgi | udpgal_g + ksi_pre8_g <-> h_g + udp_g + ksi_pre9_g | Keratan sulfate biosynthesis | golgi | B4galt1; B4galt5 | 0.55866;0.41667 |
| G14T9g | Beta-N-acetylglucosaminylglycopeptide beta-1,4-galactosyltransferase, Golgi | udpgal_g + ksi_pre11_g <-> h_g + udp_g + ksi_pre12_g | Keratan sulfate biosynthesis | golgi | B4galt1; B4galt5 | 0.55866;0.41667 |
| G14Tg | Beta-N-acetylglucosaminylglycopeptide beta-1,4-galactosyltransferase | fn2m2masn_g + 2.0 udpgal_g <-> 2.0 udp_g + l2fn2m2masn_g + 2.0 h_g | N-Glycan Biosynthesis | golgi | B4galt1; B4galt5 | 0.55866;0.41667 |
| NAt3_1 | Sodium proton antiporter HNA is 11 | na1_c + h_e <-> h_c + na1_e | Transport, Extracellular | golgi | Slc9a2 | 0.62893 |
| NAt5 | Sodium/ammonium proton antiporter | nh4_e + na1_c <-> na1_e + nh4_c | Transport, Extracellular | golgi | Slc9a2 | 0.62893 |
| PDE1g | 3',5'-cyclic-nucleotide phosphodiesterase,Golgi | camp_g + h2o_g <-> amp_g + h_g | Nucleotides | golgi | Pde10a | 0.36765 |
| PDE4g | 3',5'-cyclic-nucleotide phosphodiesterase, Golgi | 35cgmp_g + h2o_g <-> h_g + gmp_g | Nucleotides | golgi | Pde10a | 0.36765 |
| S23T2g | Beta-galactoside alpha-2,3-sialyltransferase (core 2) | cmpacna_g + core2_g <-> ksii_core2_pre1_g + h_g + cmp_g | Keratan sulfate biosynthesis | golgi | St3gal1 | 0.54945 |
| S23T4g | Beta-galactoside alpha-2,3-sialyltransferase | cmpacna_g + ksii_core4_pre1_g <-> cmp_g + ksii_core4_pre2_g + h_g | Keratan sulfate biosynthesis | golgi | St3gal1 | 0.54945 |
| S23Tg | Beta-galactoside alpha-2,3-sialyltransferase (T antigen) | T_antigen_g + cmpacna_g <-> sT_antigen_g + h_g + cmp_g | O-Glycan Biosynthesis | golgi | St3gal1 | 0.54945 |
| UGALGTg | UDPgalactose:D-glucose 4-beta-D-galactosyltransferase, Golgi apparatus | udpgal_g + glc__D_g <-> h_g + udp_g + lcts_g | Galactose metabolism | golgi | B4galt1 | 0.55866 |

| | | | | | | |
|---|---|---|---|---|---|---|
| 5HOXINDACTOXm | 5-Hydroxyindoleacetaldehyde:NAD+ oxidoreductase (mito) | nad_m + 5hoxindact_m + h2o_m <-> 2.0 h_m + nadh_m + 5hoxindoa_m | Tryptophan metabolism | mitochondria | Aldh2 | 1.67 |
| AGPRim | N acetyl g glutamyl phosphate reductase irreversible mitochondrial | acg5p_m + h_m + nadph_m <-> acg5sa_m + nadp_m + pi_m | Urea cycle/amino group metabolism | mitochondria | Aldh18a1 | 0.4065 |
| ALASm | 5 aminolevulinate synthase | gly_m + h_m + succoa_m <-> 5aop_m + co2_m + coa_m | Glycine, Serine, and Threonine Metabolism | mitochondria | Gcat | 1.82 |
| ALATA_L | L-alanine transaminase | akg_c + ala__L_c <-> glu__L_c + pyr_c | Glutamate metabolism | mitochondria | Gpt2 | 1.51 |
| ALDD20xm | Aldehyde dehydrogenase indole 3 acetaldehyde NAD mitochondrial | h2o_m + id3acald_m + nad_m <-> 2.0 h_m + ind3ac_m + nadh_m | Tryptophan metabolism | mitochondria | Aldh2 | 1.67 |
| ALDD2xm | Aldehyde dehydrogenase acetylaldehyde NAD mitochondrial | acald_m + h2o_m + nad_m <-> ac_m + 2.0 h_m + nadh_m | Glycolysis/Gluconeogenesis | mitochondria | Aldh2 | 1.67 |
| BAMPPALDOXm | Beta-Aminopropion aldehyde:NAD+ oxidoreductase (m) | nad_m + h2o_m + bamppald_m <-> nadh_m + ala_B_m + 2.0 h_m | beta-Alanine metabolism | mitochondria | Aldh2 | 1.67 |
| BDHm | 3-Hydroxybutanoate:NAD+ oxidoreductase, mitochondria | nad_m + bhb_m <-> h_m + acac_m + nadh_m | Butanoate Metabolism | mitochondria | Bdh1 | 0.67114 |
| CK | ATP Creatine kinase | creat_m + atp_m <-> adp_m + pcreat_m | Urea cycle/amino group metabolism | mitochondria | Ckmt2 | 0.51813 |
| CYANSTm | Cyanide sulfurtransferase, mitochondrial | cyan_m + tsul_m <-> tcynt_m + so3_m + h_m | Cysteine Metabolism | mitochondria | Tst | 1.9 |
| G5SDym | Glutamate-5-semialdehyde dehydrogenase, mitochondria | h_m + nadph_m + glu5p_m <-> nadp_m + pi_m + glu5sa_m | Urea cycle/amino group metabolism | mitochondria | Aldh18a1 | 0.4065 |
| GCALDDm | Glycolaldehyde dehydrogenase, mitochondrial | nad_m + h2o_m + gcald_m <-> nadh_m + 2.0 h_m + glyclt_m | Glyoxylate and Dicarboxylate Metabolism | mitochondria | Aldh2;Aldh18a1 | 1.67;0.4065 |
| GLACOm | D-Glucuronolactone:NAD+ oxidoreductase, mitochondrial | glac_m + 2.0 h2o_m + nad_m <-> glcr_m + 3.0 h_m + nadh_m | Ascorbate and Aldarate Metabolism | mitochondria | Aldh2 | 1.67 |

| | | | | | | |
|---|---|---|---|---|---|---|
| GLU5Km | Glutamate 5-kinase (m) | glu__L_m + atp_m <-> glu5p_m + adp_m | Urea cycle/amino group metabolism | mitochondria | Aldh18a1 | 0.4065 |
| GLUTCOADHm | Glutaryl-CoA dehydrogenase (mitochondria) | fad_m + h_m + glutcoa_m <-> fadh2_m + co2_m + b2coa_m | Tryptophan metabolism | mitochondria | Gcdh | 1.6 |
| GLYATm | Glycine C-acetyltransferase | gly_m + accoa_m <-> 2aobut_m + coa_m | Glycine, Serine, and Threonine Metabolism | mitochondria | Gcat | 1.82 |
| IMACTD_m | Imidazole acetaldeyde dehydrogenase (mito) | im4act_m + h2o_m + nad_m <-> im4ac_m + nadh_m + 2.0 h_m | Histidine Metabolism | mitochondria | Aldh2 | 1.67 |
| LCADi_Dm | Lactaldehyde dehydrogenase, mitochondrial | nad_m + lald__D_m + h2o_m <-> nadh_m + 2.0 h_m + lac__D_m | Pyruvate Metabolism | mitochondria | Aldh2;Aldh18a1 | 1.67;0.4065 |
| LCADm | Lactaldehyde dehydrogenase, mitochondrial | nad_m + h2o_m + lald__L_m <-> 2.0 h_m + lac__L_m + nadh_m | Pyruvate Metabolism | mitochondria | Aldh2;Aldh18a1 | 1.67;0.4065 |
| NABTNOm | N4-Acetylaminobutanal:NAD+ oxidoreductase (m) | nad_m + h2o_m + n4abutn_m <-> nadh_m + 2.0 h_m + 4aabutn_m | Arginine and Proline Metabolism | mitochondria | Aldh2 | 1.67 |
| NADPNe | NADP nucleosidase,extracellular | nadp_e + h2o_e <-> adprbp_e + ncam_e + h_e | NAD Metabolism | mitochondria | Cd38 | 0.54054 |
| OIVD1m | 2-oxoisovalerate dehydrogenase (acylating; 4-methyl-2-oxopentaoate), mitochondrial | nad_m + 4mop_m + coa_m <-> ivcoa_m + co2_m + nadh_m | Valine, Leucine, and Isoleucine Metabolism | mitochondria | Bckdhb | 1.94 |
| OIVD2m | 2-oxoisovalerate dehydrogenase (acylating; 3-methyl-2-oxobutanoate), mitochondrial | nad_m + coa_m + 3mob_m <-> co2_m + nadh_m + ibcoa_m | Valine, Leucine, and Isoleucine Metabolism | mitochondria | Bckdhb | 1.94 |
| OIVD3m | 2-oxoisovalerate dehydrogenase (acylating; 3-methyl-2-oxopentanoate), mitochondrial | coa_m + 3mop_m + nad_m <-> co2_m + 2mbcoa_m + nadh_m | Valine, Leucine, and Isoleucine Metabolism | mitochondria | Bckdhb | 1.94 |
| P45027A11m | 5-beta-cholestane-3-alpha,7-alpha,12-alpha-triol 27-hydroxylase | nadph_m + o2_m + h_m + xoltriol_m <-> xoltetrol_m + nadp_m + h2o_m | Bile Acid Biosynthesis | mitochondria | Cyp27a1 | 1.59 |
| P45027A12m | 5-beta-cholestane-3-alpha,7-alpha,12-alpha-triol 27-hydroxylase | xoltetrol_m + nadp_m <-> h_m + thcholst_m + nadph_m | Bile Acid Biosynthesis | mitochondria | Cyp27a1 | 1.59 |

| | | | | | | |
|---|---|---|---|---|---|---|
| P45027A13m | 5-beta-cholestane-3-alpha,7-alpha,12-alpha-triol 27-hydroxylase | nadph_m + o2_m + thcholst_m <-> nadp_m + thcholstoic_m + h2o_m | Bile Acid Biosynthesis | mitochondria | Cyp27a1 | 1.59 |
| P45027A14m | 5-beta-cytochrome P450, family 27, subfamily A, polypeptide 1 | nadph_m + o2_m + h_m + xol7ah2_m <-> nadp_m + h2o_m + xol7ah3_m | Bile Acid Biosynthesis | mitochondria | Cyp27a1 | 1.59 |
| P45027A15m | 5-beta-cytochrome P450, family 27, subfamily A, polypeptide 1 | nadph_m + o2_m + h_m + xol7ah3_m <-> nadp_m + 2.0 h2o_m + xol7ah2al_m | Bile Acid Biosynthesis | mitochondria | Cyp27a1 | 1.59 |
| P45027A16m | Cytochrome P450 27 | o2_m + nadph_m + xol7ah2al_m <-> dhcholestanate_m + nadp_m + h2o_m | Bile Acid Biosynthesis | mitochondria | Cyp27a1 | 1.59 |
| P45027A1m | Cytochrome P450 27 | chsterol_m + nadph_m + o2_m + h_m <-> xol27oh_m + nadp_m + h2o_m | Bile Acid Biosynthesis | mitochondria | Cyp27a1 | 1.59 |
| PYLALDOXm | Perillyl aldehyde:NAD+ oxidoreductase (m) | pylald_m + nad_m + h2o_m <-> nadh_m + 2.0 h_m + peracd_m | Limonene and pinene degradation | mitochondria | Aldh2 | 1.67 |
| VITD2Hm | Vitamin D-25-hydroxylase (D2) | h_m + nadph_m + vitd2_m + o2_m <-> h2o_m + nadp_m + 25hvitd2_m | Vitamin D | mitochondria | Cyp27a1 | 1.59 |
| VITD3Hm | Vitamin D-25-hydroxylase (D3) | o2_m + nadph_m + h_m + vitd3_m <-> nadp_m + h2o_m + 25hvitd3_m | Vitamin D | mitochondria | Cyp27a1 | 1.59 |
| LYSMTF1n | Histone-lysine N-methyltransferase, nuclear | amet_n + peplys_n <-> Nmelys_n + ahcys_n | Lysine Metabolism | nucleus | Dot1l | 0.58824 |
| LYSMTF2n | Histone-lysine N-methyltransferase, nuclear | Nmelys_n + amet_n <-> Ndmelys_n + ahcys_n | Lysine Metabolism | nucleus | Dot1l | 0.58824 |
| LYSMTF3n | Histone-lysine N-methyltransferase, nuclear | amet_n + Ndmelys_n <-> ahcys_n + Ntmelys_n | Lysine Metabolism | nucleus | Dot1l | 0.58824 |
| PTHPSn | 6-pyruvoyltetrahydropterin synthase, nuclear | ahdt_n <-> 6pthp_n + pppi_n | Tetrahydrobiopterin | nucleus | Pts | 1.64 |
| XANDp | Xanthine dehydrogenase, peroxisomal | h2o_x + nad_x + xan_x <-> urate_x + h_x + nadh_x | Purine Catabolism | peroxisome | Xdh | 1.77 |
| XAO2x | Xanthine oxidase | hxan_x + o2_x + h2o_x <-> xan_x + h2o2_x | Purine Catabolism | peroxisome | Xdh | 1.77 |
| XAOx | Xanthine oxidase,peroxisomal | o2_x + h2o_x + xan_x <-> urate_x + h2o2_x | Purine Catabolism | peroxisome | Xdh | 1.77 |
| PROAKGOX1r | L-Proline,2-oxoglutarate:oxygen oxidoreductase (4-hydroxylating) (ER) | pro__L_r + o2_r + akg_r <-> co2_r + succ_r + 4hpro_LT_r | Arginine and Proline Metabolism | ribosome | P4ha1 | 0.64516 |
